# Chronic stress reduces behavioural flexibility and autonomic adaptation when aversive outcomes are controllable

**DOI:** 10.64898/2026.11.19.767963

**Authors:** Daniyal Rajput, Kim Felmingham, Chin-Hsuan Sophie Lin, Marta I. Garrido

## Abstract

Chronic stress is associated with altered adaptation to threat, yet its effects on aversive learning under varying threat predictability and controllability remain unclear. Thirty participants with chronic stress and thirty healthy controls completed a task to learn how well visual cues predicted upcoming electric shocks (predictability) and whether correct predictions of shocks could reduce shock intensity (controllability). We found that, for controllable conditions, accuracy was similar across varying degrees of predictability in the chronic stress group, whereas accuracy in the healthy control group was highest in highly predictable blocks. The chronic stress group responded more slowly than controls and their skin conductance responses were generally lower and indistinguishable across predictability and controllability. We used a computational model to estimate subjective uncertainty and related these estimates to behavioural responses and skin conductance. Modelling showed that controllability shaped uncertainty differently across groups. In chronic stress, with control, subjective stress reflected uncertainty about likely outcomes, reaction time reflected changing outcome probabilities, and skin conductance reflected unexpected outcomes. Without control, subjective stress reflected unexpected outcomes, reaction time uncertainty about likely outcomes, and skin conductance both unexpected outcomes and outcome uncertainty. In healthy controls, subjective stress reflected changing outcome probabilities with control and outcome uncertainty without control, whereas skin conductance showed the reverse pattern, and reaction time reflected likely outcomes only under control. Together, these findings indicate less adaptation in chronic stress, reflected in reduced modulation of reaction time by controllability and skin conductance by predictability and controllability, alongside altered uncertainty associations across response domains.

## Introduction

Chronic stress is associated with sustained alterations in central and autonomic nervous system function and endocrine stress-response systems [1–3]. These alterations are accompanied by changes in physiological stress responses and in neural systems involved in stress regulation [2–6]. Animal and human studies further show that chronic stress is associated with reduced flexibility in adapting behaviour when environmental conditions change [6–8]. However, it remains unclear how these biological alterations contribute to behavioural and physiological changes during aversive learning.

These alterations may be particularly relevant when individuals with chronic stress are exposed to acute stressors. Responding to acute stress requires environmental information to evaluate threats and adjust behavioural and autonomic responses accordingly [9, 10]. This involves learning whether threat can be anticipated (predictability) and whether outcomes can be influenced by behaviour (controllability). Chronic stress may therefore alter both the magnitude of stress responses and how environmental information is used to guide those responses [11, 12]. Distinguishing these possibilities is important because reduced adaptation could arise from difficulty using reliable predictive information, reduced sensitivity to behavioural control, or changes in how these sources of information are combined. Addressing this question therefore requires consideration of predictability and controllability together.

Predictability refers to the degree to which future outcomes can be reliably anticipated from available cues, whereas controllability refers to the extent to which behaviour can influence those outcomes. Unpredictable threat is associated with sustained anxiety and elevated physiological arousal relative to predictable threat [10, 13], whereas uncontrollable aversive events are associated with behavioural passivity and impaired fear extinction [14], alongside dysregulation of neuroendocrine stress systems [2]. Despite their shared importance in shaping responses to threat, predictability and controllability have largely been examined separately and primarily in healthy individuals. It therefore remains unclear how these factors jointly influence aversive learning in individuals with chronic stress, particularly when behaviour can be used to reduce the aversive consequences of an anticipated outcome.

Hierarchical Bayesian models provide a computational account of how environmental structure is inferred from experience during aversive learning [15]. Within these models, belief updating is characterised by distinct forms of uncertainty: 1) surprise quantifies the discrepancy between expected and observed outcomes at any given moment; 2) belief uncertainty reflects uncertainty in current expectations about outcome probabilities, and 3) volatility represents the inferred rate of environmental change [16–19]. Together, these quantities determine how strongly new observations update prior beliefs. In healthy individuals, subjective stress ratings, skin conductance, and pupil dilation scale with inferred outcome uncertainty rather than objective shock frequency during probabilistic aversive learning, showing the relevance of internal estimates of uncertainty over behaviour [15]. Acute tress has been associated with reduced flexibility in updating aversive value when outcome contingencies change [20], suggesting that acute stress can alter the computational processes through which new information is incorporated into existing expectations.

A growing body of evidence suggests that chronic stress may alter multiple aspects of those computational processes. Theoretical accounts of stress-related psychopathology suggest that persistent stress may bias the processing of new information, reducing flexibility in belief updating and adaptation to changing outcomes [21]. Consistent with this theory, chronic stress has been associated with reduced behavioural flexibility, diminished fear extinction and threat-safety discrimination [12, 22, 23]. Chronic stress has also been associated with difficulties using action– outcome relationships to guide behaviour, making it harder to avert adverse outcomes [22.25]. There is a need to disentangle how the estimation of different forms of uncertainty changes in chronic stress. Given that adapting to changes in the environment requires individuals to learn whether their actions can influence outcomes, interactions between the perceived availability of control over aversive outcomes and those computational processes under chronic stress also warrant investigation. Importantly, subjective experience, behavioural performance, and autonomic responses need not express the same computational quantity. Examining these domains in parallel can help determine whether chronic stress is associated with general alterations in uncertainty processing or whether these alterations differ across response measures and controllability conditions.

The present study addressed this issue by comparing adults with chronic stress and healthy controls using a probabilistic aversive-learning task in which predictability varied across three probability levels and controllability was manipulated across sessions. Trial-wise learning was modelled using the Volatile Kalman Filter (VKF), which estimates surprise, belief uncertainty, environmental volatility, and learning rate [19]. We tested whether chronic stress was associated with differences in behavioural and autonomic responses across predictability and controllability conditions, and whether the computational quantities associated with subjective stress, reaction time, and skin conductance differed between groups. Based on evidence that chronic stress reduces flexibility and alters belief updating, we hypothesised weaker behavioural and autonomic adjustment to predictability and controllability in chronic stress than in healthy controls, and that computational quantities associated with stress ratings, reaction time, and skin conductance would differ between groups and by controllability.

## Methods and Materials

### Participants

Sixty adults aged 18 to 40 years were recruited, comprising 30 individuals with chronic stress and 30 healthy controls. Chronic stress was defined using the Trier Inventory for Chronic Stress, Short Form (TICS-9) [26,43]. Participants entered the chronic stress group if they scored 27 or higher and reported persistent stress for at least three months [43]. Eligibility criteria included no history of neurological, psychiatric, or cardiovascular disorders and no current smoking or substance use. Before testing, participants completed the Beck Depression Inventory [27], Beck Anxiety Inventory [28], and Intolerance of Uncertainty Scale [29], including prospective (desire for predictability) and inhibitory (difficulty acting under uncertainty) subscale scores. Further details on questionnaire scoring are provided in the Supplementary Methods. All participants provided written informed consent. The study was approved by the Human Research Ethics Committee of the University of Melbourne (025-31388-64207-4).

### Task and Design

Details of the experimental procedure have been reported previously [30]. In brief, participants completed a probabilistic aversive-learning task with electric shock as the aversive outcome (Fig. 1). Group (healthy control, chronic stress) was a between-participant factor, and Controllability (controllable, uncontrollable) and Predictability (high, moderate, unpredictable) were within-participant factors. Each session comprised 320 trials and was preceded by task instructions and practice. Full procedural details are provided in the Supplemental Methods.

**Figure 1.**
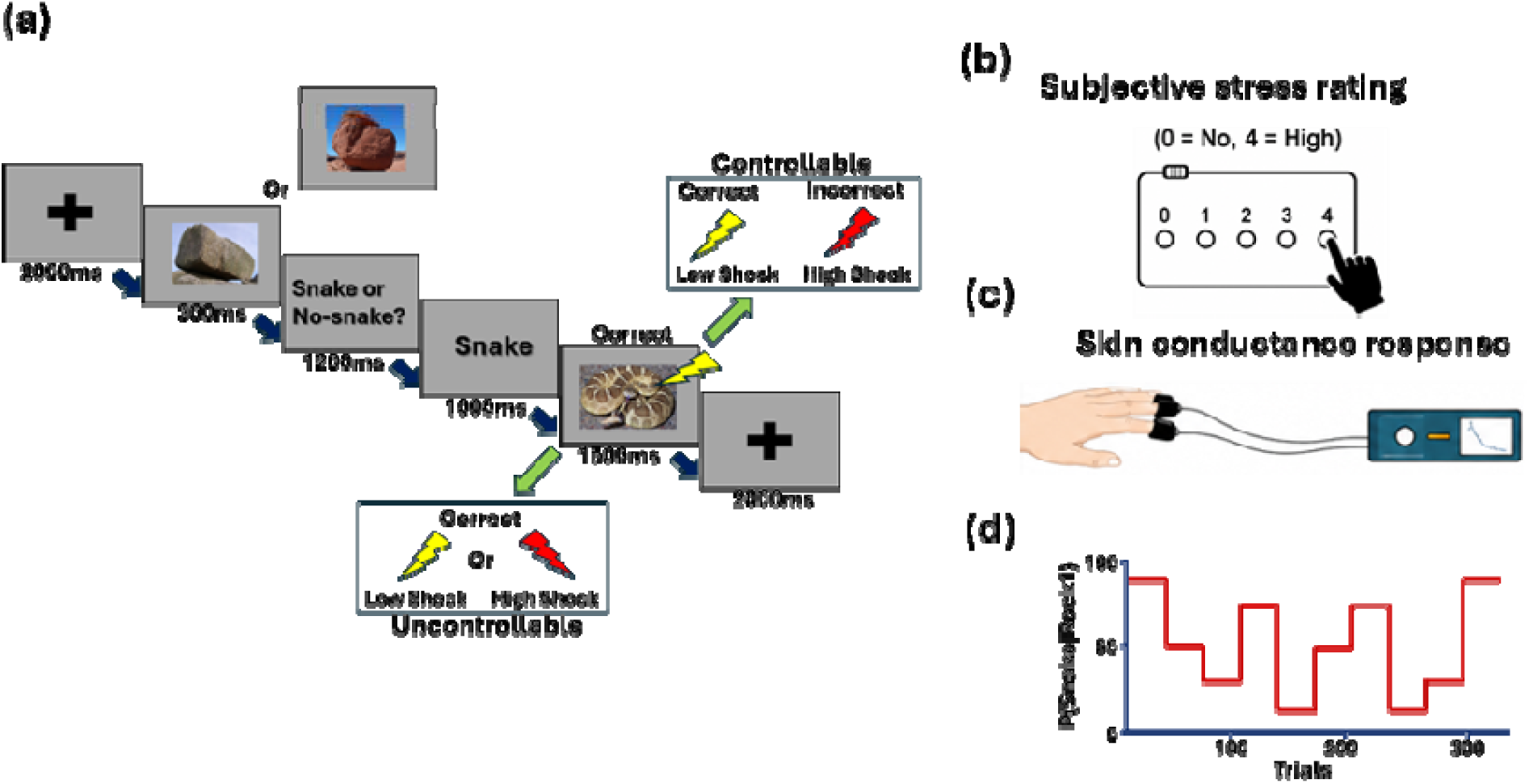
Schematic overview of the probabilistic aversive-learning task. **(a)** On each trial, participants viewed on of two rock cues, predicted whether a snake would appear, and received feedback indicating whether the snake was present or absent. Snake-present outcomes were accompanied by an electric shock. In the controllable condition, cor ect predictions reduced shock intensity, whereas incorrect predictions resulted in a high-intensity shock. In the uncontrollable condition, shock intensity was independent of prediction accuracy. **(b)** Participants intermittently rated their subjective stress on a five-point scale. **(c)** Skin conductance was recorded continuously from the non-dominant hand. **(d)** Cue– outcome probabilities varied across 10 unsignalled blocks, including highly predictable, moderately predictable, and unpredictable blocks.

On each trial, participants predicted whether a snake outcome would occur after one of two visual cues. Snake outcomes were paired with electric shock. Subjective stress ratings were obtained approximately every four to six trials (65 ratings per session) on a five-point scale, and skin conductance was recorded continuously from the nondominant hand at 1000 Hz.

Participants completed controllable and uncontrollable sessions on separate days. In the controllable condition, correct prediction of a snake outcome resulted in a low-intensity shock and an incorrect prediction in a high-intensity shock, allowing accurate prediction to reduce shock intensity. In the uncontrollable condition, shock intensity was independent of prediction accuracy. Predictability was defined by cue-outcome probabilities of 90/10% (high), 70/30% (moderate), and 50/50% (unpredictable), varying across 10 unsignalled blocks. Participants were informed that cue-outcome probabilities could change during the task.

### Shock Calibration and Physiological Measures

Shock intensity was individually calibrated before each session using an established staircase procedure [31], with the low-intensity shock set at 40% of the high-intensity level. Prediction accuracy was calculated as the proportion of correct predictions across valid shock trials, and reaction time was measured from cue onset to response. Anticipatory skin conductance response was defined as the peak phasic response during the 2 seconds preceding outcome presentation. Acquisition and preprocessing details are provided in the Supplementary information.

### Statistical and Computational Analysis

Prediction accuracy was analysed using a linear mixed-effects model with Group, Controllability, Predictability, and their interactions; a 2 × 2 × 3 mixed repeated-measures analysis of variance was used as a robustness check. Reaction time and skin conductance responses were analysed using 2 × 2 × 3 mixed analyses of variance. Significant effects were followed by paired-or independent-samples t tests with Bonferroni correction.

Trial-by-trial belief updating was modelled separately for each participant and session using the VKF [19]. The model generated trial-wise estimates of surprise, belief uncertainty, environmental volatility, and learning rate. Surprise reflected the discrepancy between expected and observed outcomes, belief uncertainty reflected uncertainty in the current belief estimate, and volatility indexed the inferred rate of change in outcome contingencies. Model specification and fitting are described in the Supplemental Methods.

Participant-level general linear models related these computational quantities to subjective stress ratings, reaction times, and skin conductance responses within each controllability condition. Surprise, belief uncertainty, and volatility were entered in separate competing models; prediction accuracy was included in all models, and the preceding stress rating was additionally included for subjective ratings. Models were compared separately by response measure, group, and controllability using random-effects Bayesian model comparison, with exceedance probability identifying the best-supported model. Regression coefficients were examined for effects of group and controllability. Secondary learning-rate, volatility-update, and questionnaire analyses are reported in the Supplementary information.

## Results

### Prediction accuracy differed between groups across controllability and predictability conditions

Behavioural analyses first tested whether group differences in prediction accuracy depended on controllability and predictability (Fig. 2). A linear mixed-effects model revealed a significant main effect of Group, F(1,305.73)=4.63, p=0.032, with higher overall prediction accuracy in the chronic stress group than in healthy controls. together with significant Group × Predictability, F(2,285.19)=20.57, p<0.001, Controllability × Predictability, F(2,284.86)=5.41, p=0.005, and Group × Controllability × Predictability interactions, F(2,284.51)=5.10, p=0.007 (Fig. 1). No significant main effects of Controllability or Predictability were observed. A mixed repeated-measures analysis of variance showed a similar pattern.

**Fig. 2.**
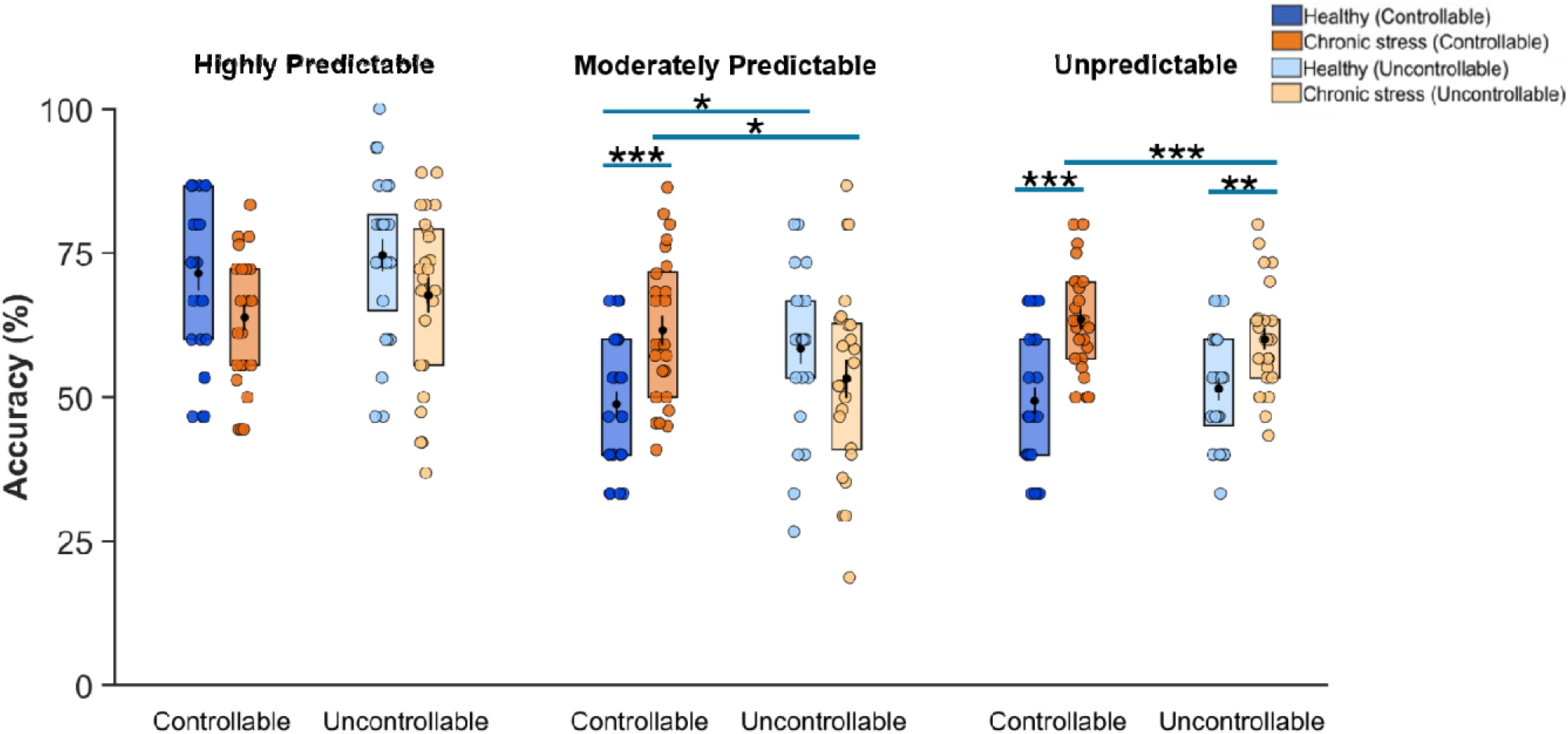
Prediction accuracy across controllability, predictability and group. Prediction accuracy, defined as the proportion of trials on which participants correctly predicted the outcome (snake) and thereby avoided high-intensity shock in the controllable condition, is shown for healthy controls (blue) and individuals with chronic stress (orange) under controllable (light) and uncontrollable (dark) conditions at high, moderate and unpredictable levels of predictability. Points represent individual participants. Boxplots show the median and interquartile range, with whiskers extending to 1.5 × the interquartile range. Linear mixed-effects modelling revealed a main effect of group, F(1,305.73) = 4.63, p = 0.032, together with group × predictability, F(2,285.19) = 20.57, p <0.001, and group × controllability × predictability, F(2,284.51) = 5.10, p = 0.007, interactions.

In the controllable condition, the chronic stress group showed higher accuracy than healthy controls at moderate predictability and under unpredictable conditions, both p<0.001, whereas no group difference was observed at high predictability. In the uncontrollable condition, the group difference was significant only when outcomes were unpredictable, with the chronic stress group exhibiting higher prediction accuracy than healthy controls, p=0.004. Controllability also affected the groups differently at moderate predictability: healthy controls were more accurate under uncontrollable than controllable conditions, p=0.039, whereas the chronic stress group showed the opposite pattern, p=0.027. The chronic stress group also showed higher accuracy under controllable than uncontrollable conditions when outcomes were unpredictable, p<0.001. Healthy controls were most accurate under high predictability in both controllability conditions, all p<0.001. In contrast, the chronic stress group showed a high-predictability advantage only under uncontrollable conditions relative to moderate predictability, p=0.007.

### Slower responses for the chronic stress group and faster responses under uncontrollable conditions for both groups

A 2 × 2 × 3 mixed analysis of variance showed a significant main effect of Group, with slower responses in the chronic stress group than healthy controls, F(1,51)=207.11, p<0.001, and a main effect of Controllability, with faster responses under uncontrollable than controllable conditions, F(1,51)=16.24, p<0.001 (Fig. 3). There was no main effect of Predictability, all Bonferroni-corrected p>.50. A Group × Controllability interaction was significant, F(1,51)=4.94, p=0.031. The chronic stress group had longer reaction times than healthy controls across all conditions, all correc ed p<0.001. In healthy controls, reaction times were longer under controllable than uncontrollable conditions at high predictability, t(26)=4.46, p<.001, and moderate predictability, t(26)=3.82, p=0.004, but not under unpredictable conditions, p=0.412. In the chronic stress group, reaction times did not differ significantly between controllability conditions at any predictability level, all corrected p>0.57.

**Fig. 3.**
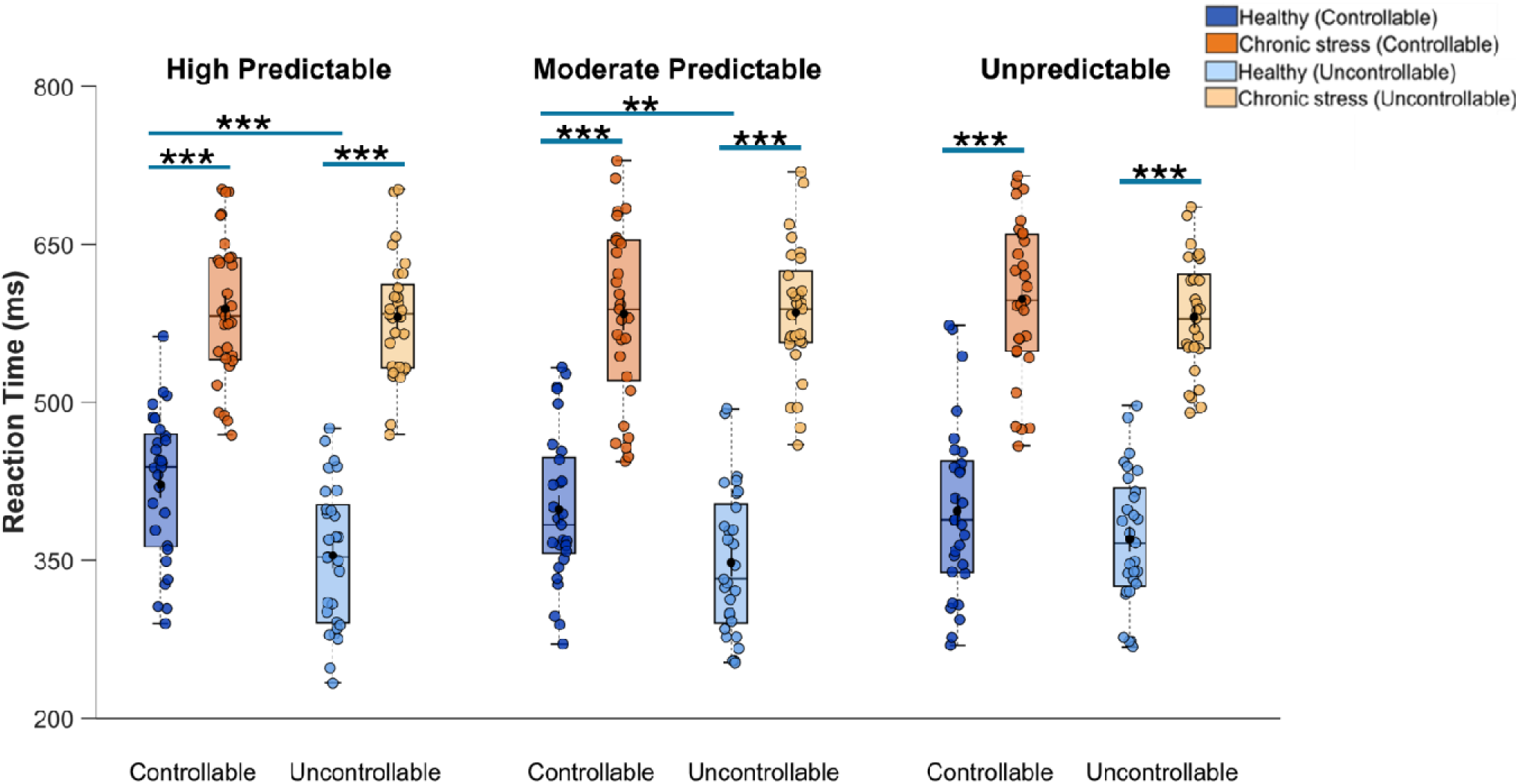
Reaction time across controllability, predictability and group. Reaction time (ms), measured as the interval between cue onset and the participant’s response, for healthy controls (blue) and individuals with chronic stress (orange) under controllable (light) and uncontrollable (dark) conditions at high, moderate and unpredictable levels of predictability. Points denote individual participants and error bars represent the mean ± SEM. A 2 × 2 × 3 mixed analysis of variance revealed main effects of group, F(1,51) = 207.11, p < 0.001, and controllability, F(1,51) = 16.24, p < 0.001, but no effect of predictability (all p > 0.50). A group × controllability interaction was observed, F(1,51) = 4.94, p = 0.031, indicating slower responses in the chronic stress group and condition-dependent effects of controllability.

### Skin conductance responses showed reduced condition-dependent differentiation in chronic stress

Skin conductance responses differed between groups and varied across controllability and predictability (Fig. 4). Significant main effects were observed for Group, F(1,50)=22.47, p<0.001, higher skin conductance responses in healthy controls than in the chronic stress group. A significant main effect of controllability was found, F(1,50)=55.02, p<0.001, with higher responses under controllable than uncontrollable conditions. Predictability also showed a significant main effect, F(2,100)=73.83, p<0.001, with responses highest under unpredictable conditions, followed by moderate and high predictability. Significant interactions were also observed for Group × Controllability, F(1,50)=39.84, p<0.001, Group × Predictability, F(2,100)=58.96, p<0.001, and Group × Controllability × Predictability interactions, F(2,100)=61.02, p<0.001. In healthy controls, skin conductance responses were lower under uncontrollable than controllable conditions at high predictability, t(25)=13.11, p<0.001, but higher under uncontrollable than controllable conditions in unpredictable blocks, t(25)=3.07, p=0.015, with no difference at moderate predictability. In the chronic stress group, skin conductance responses did not differ across controllability or predictability, all p>0.05. Between groups, responses were higher in healthy controls under controllable conditions at all predictability levels, all corrected p<0.001. Under uncontrollable conditions, responses were lower in healthy controls at high predictability, t(50)=8.22, p<0.001, but higher at moderate and unpredictable levels, both corrected p<0.001.

**Fig. 4.**
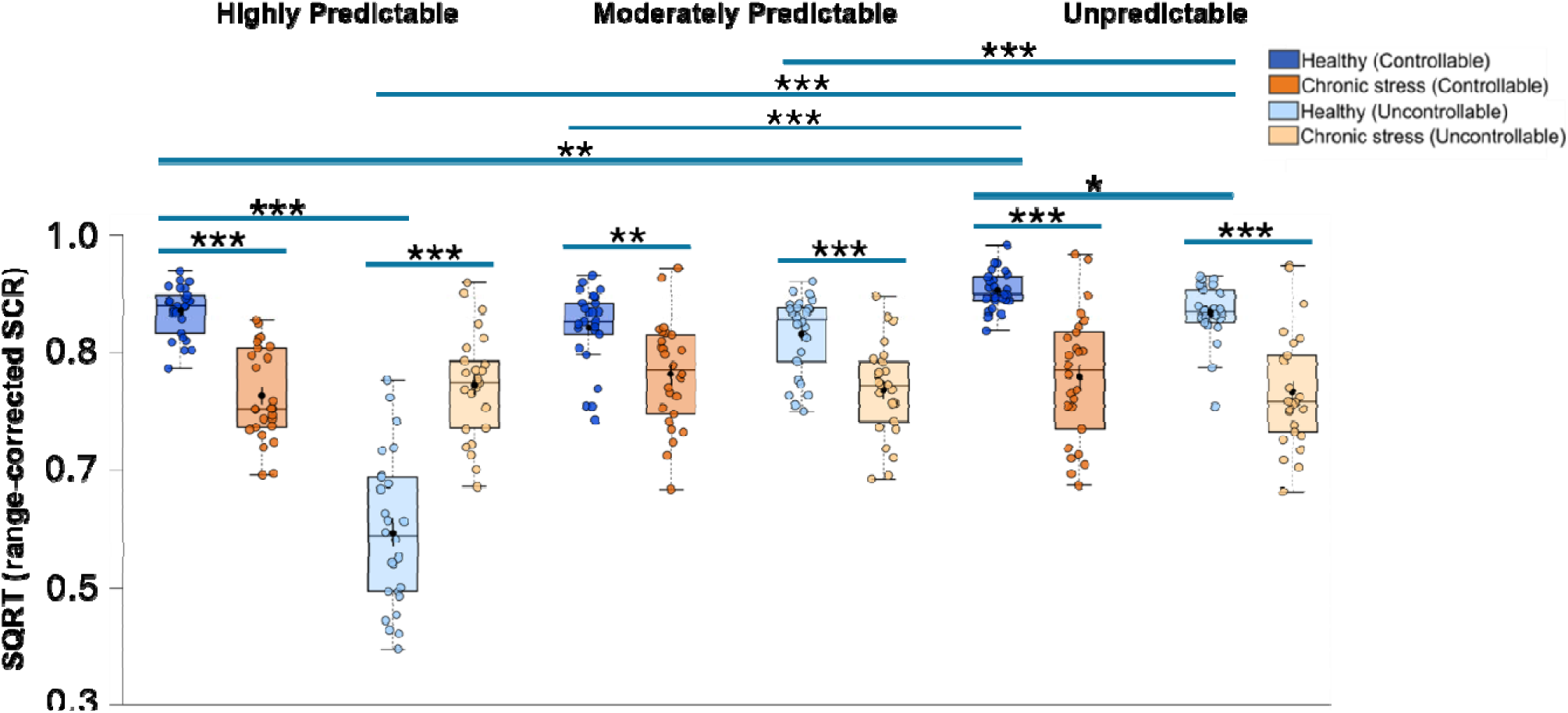
Skin conductance responses across controllability and predictability in healthy controls and individuals with chronic stress. Range-corrected skin conductance responses (SCR) for healthy controls (blue) and individuals with chronic stress (orange) under controllable (light) and uncontrollable (dark) conditions at high, moderate and unpredictable levels of predictability. Points represent individual participants; error bars represent the mean ± SEM. A 2 × 2 × 3 mixed analysis of variance revealed main effects of group, F(1,50) = 22.47, p <0.001, controllability, F(1,50) = 55.02, p <0.001, and predictability, F(2,100) = 73.83, p <0.001, together with a group × controllability × predictability interaction, F(2,100) = 61.02, p <0.001.

### Computational model preference differed between groups and controllability conditions

The computational quantity best accounting for subjective stress ratings, reaction times, and skin conductance responses differed between groups and controllability conditions (Fig. 5). For subjective stress ratings under controllable conditions, volatility showed the highest exceedance probability in healthy controls (1.000), whereas belief uncertainty showed the highest exceedance probability in the chronic stress group (0.918). Under uncontrollable conditions, belief uncertainty best explained stress ratings in healthy controls (0.943), whereas surprise showed the highest exceedance probability in chronic stress (0.545).

**Fig. 5.**
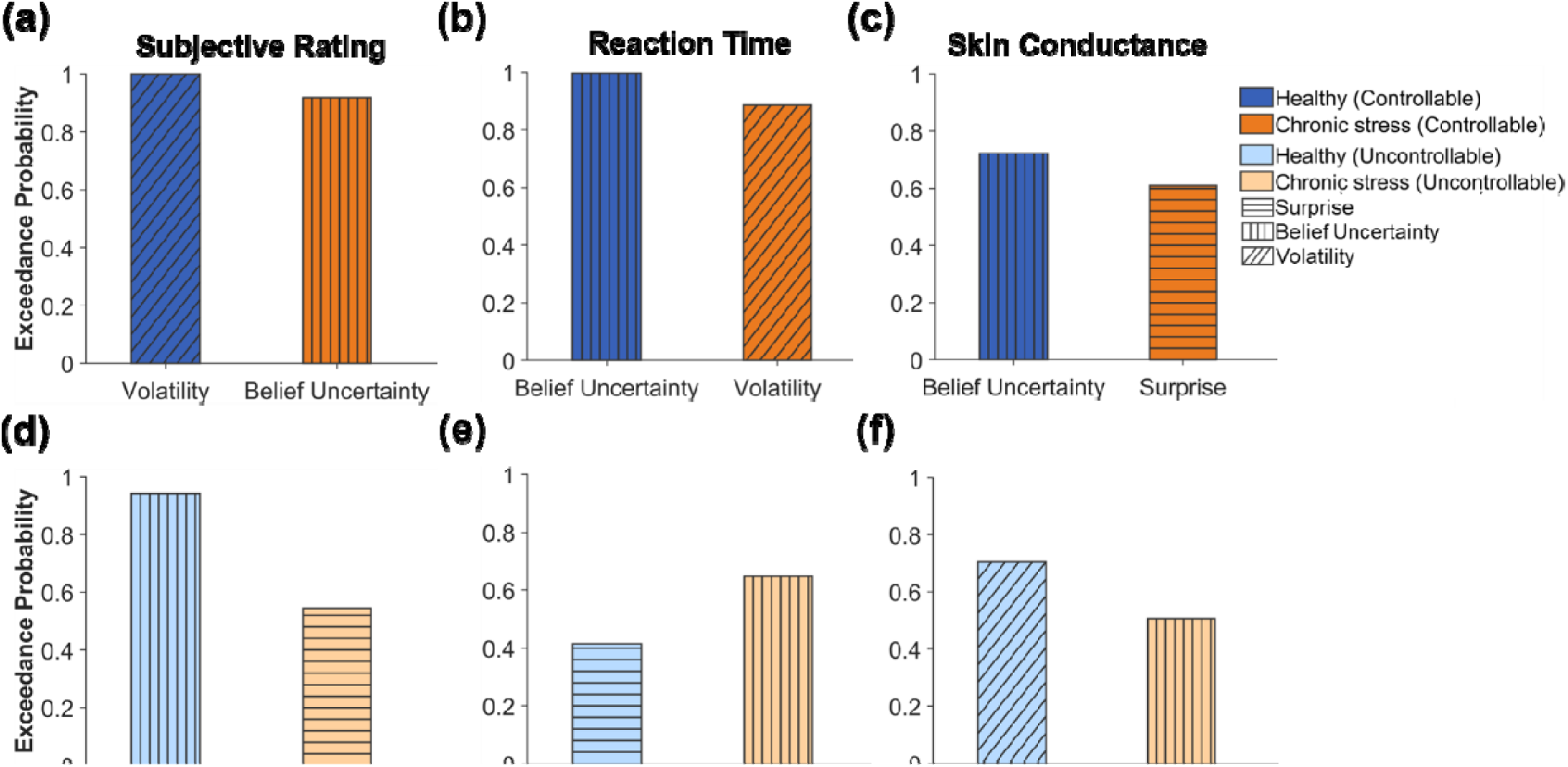
Bayesian model comparison for subjective ratings, reaction time, and skin conductance responses. Exceedance probabilities from Bayesian model comparison for surprise, belief uncertainty, and volatility in healthy controls (blue) and individuals with chronic stress (orange). Vertical, horizontal, and diagonal stripes denote belief uncertainty, surprise, and volatility, respectively. Under controllable conditions: (a) subjective ratings was best predicted by volatility in healthy participants, whereas belief uncertainty best explained subjective ratings in the chronic stress group; (b) reaction time was best explained by belief uncertainty in healthy participants, whereas volatility provided the strongest model evidence in the chronic stress group, and (c) skin conductance responses were best predicted by belief uncertainty in healthy participants, whereas surprise best explained skin conductance in the chronic stress group. Under uncontrollable conditions: (d) subjective ratings were best explained by belief uncertainty in healthy participants, whereas surprise exhibited the highest exceedance probability in the chronic stress group; (e) for reaction time, no single model clearly dominated in healthy participants, whereas belief uncertainty remained the best model in the chronic stress group; and (f) for skin conductance responses, volatility demonstrated the highest exceedance probability in healthy participants, whereas belief uncertainty showed the strongest model evidence in the chronic stress group.

For reaction times (RT), the pattern essentially reversed between groups. Specifically, under controllable conditions, belief uncertainty best explained RT in healthy controls (0.997), whereas volatility provided the strongest evidence in chronic stress (0.886). Under uncontrollable conditions, no single model predominated in healthy controls, with comparable support for surprise (0.415) and volatility (0.399), whereas belief uncertainty best predicted RT in chronic stress (0.648).

Skin conductance under controllable conditions was best predicted by belief uncertainty in healthy controls (0.722), whereas in chronic stress this was best predicted by surprise (0.610). Under uncontrollable conditions, volatility best explained skin conductance in healthy controls (0.705), whereas belief uncertainty and surprise models showed comparable performance in chronic stress (0.506 and 0.494). Thus, model preference differed between groups across all three response domains under controllable conditions, whereas the pattern under uncontrollable conditions was less consistently differentiated.

### Controllability-dependent contributions of trial-wise predictors differed between groups

Bayesian model comparison identified the best model for each response by group and controllability condition (Fig. 5). To characterise these model effects further, we then examined participant-specific regression coefficients to quantify the direction and magnitude of the relationship between each predictor and the corresponding response, and to determine whether these predictor–response relationships differed by group and controllability. Each model included prediction accuracy and one computational quantity, with subjective-rating models additionally including the preceding rating to account for dependence between successive ratings. Coefficients from the best-fitting model were examined, with models showing comparable evidence considered when no model clearly predominated. Figure 6 shows belief-uncertainty and volatility coefficients, with all models reported in Supplementary Fig. S1. The preceding-rating coefficient exceeded the prediction-accuracy coefficient. In the belief-uncertainty model, Controllability × Predictor, F(2,72)=4.66, p=0.013, and Group × Controllability × Predictor interactions, F(2,72) =3.86, p=0.025, were significant. The surprise model showed Group × Predictor, F(2,84)=3.35, p=0.040, and Controllability × Predictor interactions, F(2,84)=3.84, p=0.025. The volatility model showed a Controllability × Predictor interaction, F(2,76)=4.43, p=0.015, with no group interaction. Groups differed in the prediction-accuracy coefficient under controllable conditions, t(36)=3.09, p=0.024, and previous-rating coefficient under uncontrollable conditions, t(36)=3.62, p=0.006. For reaction time, the belief-uncertainty showed a significant predictor effect, F(1,46) = 9.29, p = 0.004, with no predictor interactions, all p > 0.05. Volatility showed Group × Controllability × Predictor interaction, F(1,48) = 9.61, p = 0.003, and a predictor effects, F(1,48) = 9.83, p = 0.003. Volatility coefficient differed by controllability in healthy controls in healthy controls, t(24) = 3.28, Bonferroni-corrected p = 0.006, but not in the chronic stress. stress. For skin conductance, volatility showed a main effect of Controllability, F(1,44)=14.09, p<0.001, and a Group × Controllability interaction, F(1,44)=19.35, p<0.001. Prediction-accuracy, t(22)=2.55, p=0.018, and volatility coefficients, t(22)=3.94, p<0.001, differed by controllability only in healthy controls. The belief-uncertainty model showed a Group × Controllability interaction, F(1,42)=7.19, p=0.010, with groups differing under controllable conditions, t(42)=3.19, p=0.004.

**Fig. 6.**
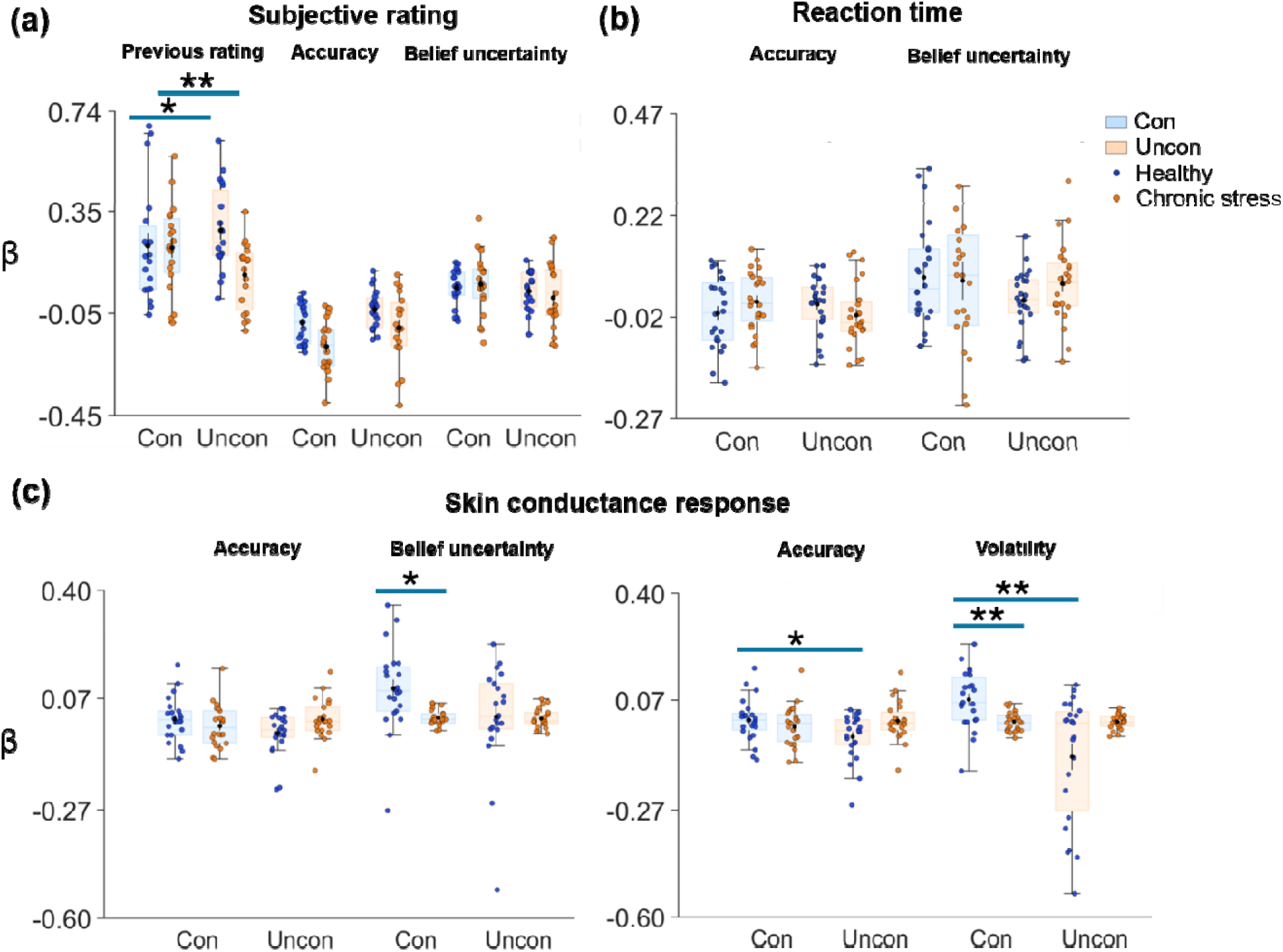
Selected regression coefficients showing differences in predictor contributions across groups and controllability conditions. Bayesian model comparison (Fig. 5) identified which computational models best explained each response. Here, regression coefficients from selected models are examined to show the direction and magnitude of each predictor’s relationship with the response, and whether these relationships differed by group and controllability. Each regression model included prediction accuracy and one computational quantity, with the preceding stress rating additionally included for subjective ratings. Regression coefficients are shown for the belief-uncertainty model of subjective ratings (a), the belief-uncertainty model of reaction time (b), and the belief-uncertainty and volatility models of skin conductance responses (c). For skin conductance, both models are shown because model preference differed across groups and controllability conditions in the Bayesian model comparison (Fig. 5). Within each predictor, Con and Uncon denote controllable and uncontrollable conditions, respectively. Healthy controls are shown in blue and individuals with chronic stress in orange, with dark shades indicating the controllable condition and light shades the uncontrollable condition. Points denote individual participants. For subjective ratings (a), coefficients for the computational predictors were smaller than those for the previous rating. The belief uncertainty model showed controllability × predictor, F(2,72) = 4.66, p = 0.013, and group × controllability × predictor effects, F(2,72) = 3.86, p = 0.025, whereas the volatility and surprise models showed controllability-dependent effects. For reaction time (b), the volatility model showed a controllability × predictor effect, F(1,48) = 7.98, p = 0.007. For skin conductance responses (c), the volatility model showed a main effect of controllability, F(1,44) = 14.09, p < 0.001, and a group × controllability interaction, F(1,44) = 19.35, p <0.001, and the belief uncertainty model also showed a group × controllability interaction, F(1,42) = 7.19, p = 0.010.

### Accuracy and learning-rate coupling robust in healthy controls but limited in chronic stress

We found a marked group difference in the association between learning rate and prediction accuracy (Fig. 7). In healthy controls, higher model-derived learning rates were associated with lower prediction accuracy across all six predictability-by-controllability conditions, all p<0.009, with the strongest associations under high predictability in controllable, r(25)=-.918, p<0.001, and uncontrollable, r(25)=-.832, p<0.001, conditions. In the chronic stress group, learning rate was negatively associated with accuracy only under controllable unpredictable conditions, r(25)=-.448, p=0.022. Additional learning-rate and volatility-update-rate analyses are reported in the

**Fig. 7.**
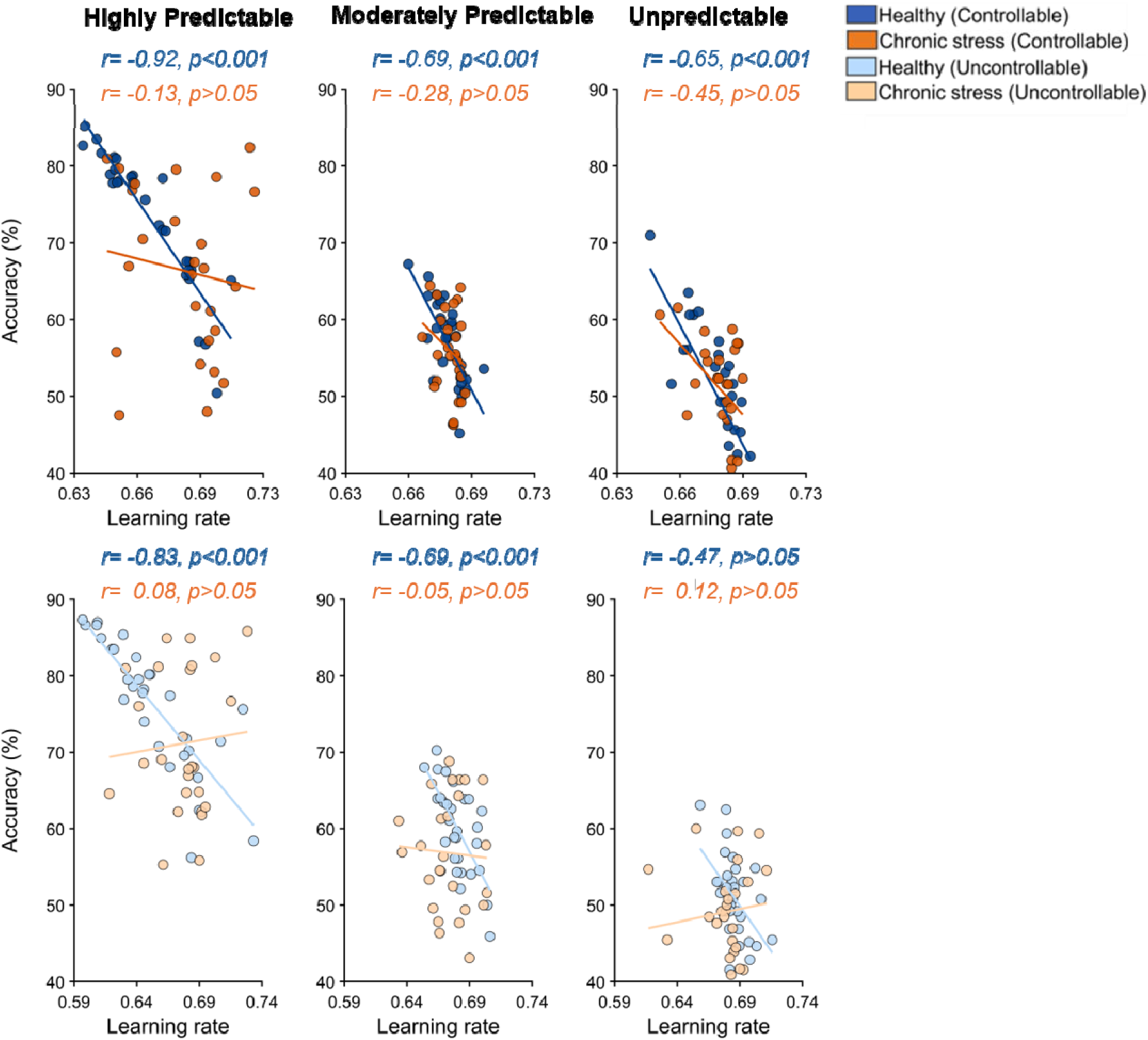
Associations between learning rate and prediction accuracy across controllability and predictability conditions in healthy controls and individuals with chronic stress. Associations between learning rate and prediction accuracy are shown across highly predictable (HP), moderately predictable (MP), and unpredictable (UP) blocks for healthy controls (blue) and individuals with chronic stress (orange) under controllable (a–c) and uncontrollable (d–f). Darker shades indicate the controllable condition and lighter shades indicate the uncontrollable condition. Points denote individual participants and lines indicate least-squares fits. In the healthy group, higher learning rates were associated with lower prediction accuracy across all predictability levels under both controllable and uncontrollable conditions (all p < 0.009), with the strongest associations observed under high predictability in controllable, r(25) = −0.918, p <0.001, and uncontrollable, r(25) = −0.832, p <0.001, conditions. In the chronic stress group, a negative association was observed only under controllable unpredictable conditions, r(25) = −0.448, p = 0.022, with no significant associations under uncontrollable conditions.

### Selective associations between self-report measures and task-derived variables

Scores on the Beck Anxiety Inventory (BAI; Fig. 8a), Beck Depression Inventory (BDI; Fig. 8b), prospective intolerance of uncertainty (IUS Factor 1; Fig. 8c), and inhibitory intolerance of uncertainty (IUS Factor 2; Fig. 8d) were higher in the chronic stress group than healthy controls, all *p*<0.001. Prospective intolerance of uncertainty reflects predictability and certainty seeking, whereas inhibitory intolerance of uncertainty reflects difficulty acting under uncertainty. No questionnaire– task associations survived Bonferroni correction in either group, all adjusted *p*>0.05. In healthy controls, uncorrected associations included BAI scores with learning rate in highly predictable blocks under the controllable condition, *r*(23)=0.51, *p*=0.009, and prediction accuracy in moderately predictable blocks, *r*(24)=−0.41, *p*=0.037. Additional associations are reported in the Supplementary Results.

**Fig. 8.**
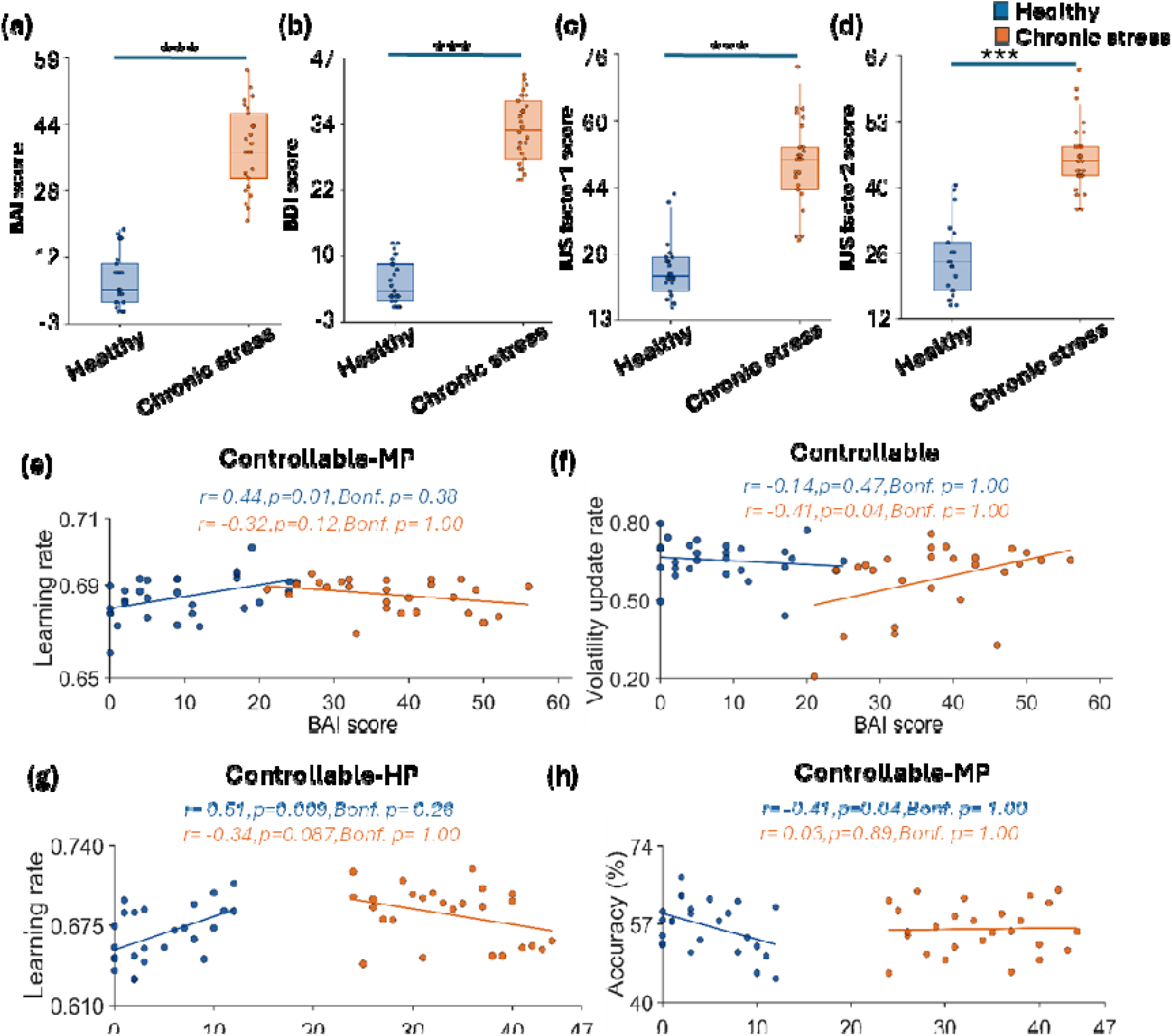
Associations between self-report measures and task-derived variables in healthy controls and individuals with chronic stress. Questionnaire scores are shown for the Beck Anxiety Inventory (BAI; a), Beck Depression Inventory (BDI; b), prospective intolerance of uncertainty (IUS Factor 1; c), and inhibitory intolerance of uncertainty (IUS Factor 2; d) in healthy controls (blue) and individuals with chronic stress (orange). Scatterplots show selected associations between questionnaire scores and task-derived measures that reached significance at the uncorrected level under the controllable condition (e–i). Points denote individual participants and lines indicate least-squares fits. Pearson’s r, uncorrected p values, and Bonferroni-adjusted p values are reported directly on each scatterplot. Questionnaire scores were higher in the chronic stress group than in healthy controls for BAI, t(44.12) = −13.75, p < 0.001, BDI, t(43.25) = −19.62, p < 0.001, and both IUS factors, both p < 0.001. Although several questionnaire–task associations reached significance before correction, none remained significant following Bonferroni correction for multiple comparisons.

## Discussion

The present findings indicated that chronic stress altered how predictability and controllability shape behavioural and autonomic responses during aversive learning, with the underlying differences explained by computational modelling. Prediction accuracy differed between groups as a function of both predictability and controllability rather than showing a general deficit in chronic stress. Reaction times were modulated by controllability in healthy controls but not in chronic stress, and skin conductance varied across predictability and controllability conditions in healthy controls but remained similar across conditions in chronic stress. Computational analyses further showed that subjective stress, reaction time, and skin conductance were predicted by different types of uncertainty across groups and controllability conditions. Together, these findings indicate altered adaptation to environmental structure in chronic stress rather than a uniform impairment in aversive learning.

Healthy controls’ prediction accuracy was supported by stable cue-outcome relationships, with the highest accuracy under highly predictable blocks and lower performance as contingencies weakened, consistent with previous work on predictability and anticipation of aversive outcomes [10, 32]. Individuals with chronic stress, however, showed higher accuracy than healthy controls when predictability was lower under controllable conditions. Rather than indicating globally better learning, the group difference emerged specifically when cue–outcome information was less reliable, and accurate predictions could reduce shock intensity. The behavioural consequences of chronic stress therefore depended on environmental structure and the relevance of prediction accuracy for the aversive outcome, consistent with evidence that stress can alter the influence of new outcome information during belief updating [33] and neural systems involved in behavioural control [1, 3].

The chronic stress group responded more slowly than healthy controls across all conditions, consistent with evidence that chronic stress alters neural systems supporting cognition and attentional control [1, 34, 35]. In healthy controls, responses were slower under controllable than uncontrollable conditions at high and moderate predictability, consistent with greater processing demands when predictions were relevant for determining shock intensity. Controllable environments promote evaluation of action-outcome contingencies, whereas uncontrollable environments can be represented independent of one’s own actions [36]. In chronic stress, reaction times did not vary with controllability. This may reflect a reduced subjective sense of control, resulting in diminished sensitivity to whether behaviour influenced shock intensity and a weaker reaction-time distinction between controllable and uncontrollable conditions. This interpretation is consistent with evidence that chronic stress is associated with less efficient cognitive control and reduced flexibility in adapting to changing task demands [11, 37], although it should be interpreted with caution because perceived controllability was not assessed directly. The absence of a predictability effect on reaction time does not imply that predictability has no behavioural effect, because accuracy and reaction time capture different aspects of performance.

Skin conductance provided converging evidence for reduced condition-dependent adaptation. In healthy controls, responses were lower under uncontrollable than controllable conditions when outcomes were highly predictable, but higher under uncontrollable conditions when outcomes were unpredictable. This is consistent with evidence that skin conductance tracks uncertainty during aversive learning [15] and is sensitive to environmental threat structure [14, 38, 39]. By contrast, SCRs in the chronic stress group were not modulated by either predictability or controllability, indicating impaired autonomic responses to variations in uncertainty and controllability, rather than a simple reduction in overall arousal. This accords with prior evidence that chronic stress is associated with blunted autonomic flexibility under changing environmental demands [5, 40].

The computational analyses clarified how these behavioural and autonomic differences were organised. In healthy controls, subjective stress was best explained by volatility when outcomes were controllable, but by belief uncertainty when outcomes were uncontrollable. Thus, subjective stress was more closely related to inferred changes in environmental contingencies when accurate predictions could influence shock intensity, but to uncertainty about current outcome probabilities when shock intensity was independent of behaviour. This is consistent with hierarchical Bayesian accounts of belief updating [17] and prior evidence linking subjective stress to computational estimates of uncertainty [15]. Belief uncertainty also best accounted for reaction time under controllable conditions in healthy controls, indicating that response latency was particularly related to uncertainty about current outcome probabilities when predictions had consequences for shock intensity.

A different organisation of computational quantities emerged in chronic stress. Under controllable conditions, subjective stress was best explained by belief uncertainty, reaction time by volatility, and skin conductance by surprise. Under uncontrollable conditions, subjective stress was best explained by surprise, reaction time by belief uncertainty, and skin conductance showed comparable evidence for belief uncertainty and surprise. Rather than a uniform shift across domains, the computational quantity associated with each response varied with controllability. The emergence of surprise as the best account of subjective stress and skin conductance in different conditions is consistent with evidence that chronic stress alters processing and adjustment following unexpected outcomes [41, 42]. The shift toward volatility in reaction time under control is consistent with evidence that stress can alter learning about changes in action–outcome contingencies [12, 24].

Comparisons of volatility regression coefficients across controllability conditions showed that the association of volatility with reaction time and skin conductance differed by controllability, particularly in healthy controls. Volatility was more strongly related to reaction time and skin conductance when accurate predictions reduced shock intensity than when shock intensity was independent of prediction accuracy, suggesting that changes in cue–outcome contingencies became more behaviourally and physiologically relevant when updated predictions could influence the aversive outcome. This is consistent with evidence that, in controllable environments, individuals incorporate the consequences of their own actions when forming predictions [36]. In chronic stress, such controllability-contingent differences in association strength between volatility estimates and behaviour were not observed, demonstrating reduced adjustment to changing environmental demands [11, 12]. For subjective ratings, the preceding-rating coefficient exceeded those for prediction accuracy and the computational quantities, indicating strong dependence on the preceding rating.

Learning-rate analyses further linked belief updating to behavioural performance. In healthy controls, higher learning rates were consistently associated with lower accuracy, with the strongest relationships under high predictability. This is consistent with normative learning accounts in which stable cue–outcome contingencies favour lower learning rates because occasional unexpected outcomes provide limited evidence that the underlying contingency has changed [16, 19]. In chronic stress, this relationship was largely absent except under controllable unpredictable conditions, suggesting that trial-wise belief updating was less significantly related to behavioural performance. Individual differences in chronic stress were instead expressed in volatility updating, with higher anxiety and inhibitory intolerance of uncertainty associated with higher volatility update rates under controllable conditions. This pattern is consistent with evidence that stress-related anxiety can increase estimates of environmental volatility and impair learning [24]. Other questionnaire associations were condition-specific and are reported in the Supplement.

A key limitation of this study is that the controllable and uncontrollable sessions were not counterbalanced, because the uncontrollable session reproduced each participant’s shock sequence from the controllable session. This compromise was chosen such that the sequence of aversive exposure was as closely matched within participants as possible. Although session order may have contributed to condition differences, the observed effects varied across groups, predictability levels, and response measures rather than showing a uniform change in the second session. This pattern suggests that the findings are unlikely to be explained solely by practice or familiarity.

In conclusion, the present findings show that predictability and controllability jointly shape aversive learning and that chronic stress alters how these factors influence subjective, behavioural, and autonomic responses. Healthy individuals showed flexible, condition-dependent responses to different forms of uncertainty, whereas chronic stress was associated with reduced differentiation across predictability and controllability conditions. Computationally, subjective stress in chronic stress was linked to belief uncertainty under controllable conditions and surprise under uncontrollable conditions, while reaction time was linked to volatility under control. These differences were most evident when outcomes were controllable, suggesting that chronic stress is associated with reduced adaptation to situations in which behaviour can influence aversive outcomes.

## Acknowledgments

We thank Katja Petrowski for providing the TICS-9 questionnaire used to assess chronic stress in this study. We also thank members of the Cognitive Neuroscience and Computational Psychiatry Lab and the Trauma, Anxiety and Stress Lab for their support and valuable discussions throughout the study. This project was funded by the 2023 Melbourne School of Psychological Sciences Research Incentive Scheme. Daniyal Rajput was supported by an Australian Government Research Training Program Scholarship, administered by the University of Melbourne.

## Data and Code Availability

The anonymised data supporting the findings are available on Figshare [https://figshare.com/s/b44df04030c89f97ed30]. Analysis code for data processing, statistical analysis, and figure generation is available at https://github.com/daniyalrajput/chronic-stress-predictability-controllability-aversive-learning_codes.git. Computational modelling code is available at https://github.com/daniyalrajput/stress_study_KF_VKF_scripts.git

## References

1. McEwen, B.S., Neurobiological and Systemic Effects of Chronic Stress. Chronic Stress (Thousand Oaks), 2017. 1.

2. McEwen, B.S., Physiology and Neurobiology of Stress and Adaptation: Central Role of the Brain. Physiological Reviews, 2007. 87(3): p. 873–904.

3. Herman, J., Neural control of chronic stress adaptation. Frontiers in Behavioral Neuroscience, 2013. **Volume** 7 **-** 2013.

4. McEwen, B.S., C. Nasca, and J.D. Gray, Stress Effects on Neuronal Structure: Hippocampus, Amygdala, and Prefrontal Cortex. Neuropsychopharmacology, 2016. 41(1): p. 3–23.

5. Wekenborg, M.K., et al., Examining reactivity patterns in burnout and other indicators of chronic stress. Psychoneuroendocrinology, 2019. 106: p. 195–205.

6. Rosenkranz, J.A., E.R. Venheim, and M. Padival, Chronic Stress Causes Amygdala Hyperexcitability in Rodents. Biological Psychiatry, 2010. 67(12): p. 1128–1136.

7. Grissom, N. and S. Bhatnagar, Habituation to repeated stress: get used to it. Neurobiol Learn Mem, 2009. 92(2): p. 215–24.

8. McEwen, B.S., et al., Mechanisms of stress in the brain. Nat Neurosci, 2015. 18(10): p. 1353–63.

9. Sandi, C., Stress and cognition. Wiley Interdiscip Rev Cogn Sci, 2013. 4(3): p. 245–261.

10. Grillon, C., et al., Anxious responses to predictable and unpredictable aversive events. Behav Neurosci, 2004. 118(5): p. 916–24.

11. Knauft, K., et al., Perceived chronic stress influences the effect of acute stress on cognitive flexibility. Scientific Reports, 2021. 11(1): p. 23629.

12. Radenbach, C., et al., The interaction of acute and chronic stress impairs model-based behavioral control. Psychoneuroendocrinology, 2015. 53: p. 268–80.

13. Robinson, O.J., et al., The effect of induced anxiety on cognition: threat of shock enhances aversive processing in healthy individuals. Cogn Affect Behav Neurosci, 2011. 11(2): p. 217–27.

14. Maier, S.F. and M.E. Seligman, Learned helplessness at fifty: Insights from neuroscience. Psychol Rev, 2016. 123(4): p. 349–67.

15. de Berker, A.O., et al., Computations of uncertainty mediate acute stress responses in humans. Nature Communications, 2016. 7(1): p. 10996.

16. Behrens, T.E., et al., Learning the value of information in an uncertain world. Nat Neurosci, 2007. 10(9): p. 1214–21.

17. Mathys, C., et al., A Bayesian Foundation for Individual Learning Under Uncertainty. Frontiers in Human Neuroscience, 2011. **volume** 5 **-** 2011.

18. Mathys, C.D., et al., Uncertainty in perception and the Hierarchical Gaussian Filter. Frontiers in Human Neuroscience, 2014. **Volume** 8 **-** 2014.

19. Piray, P. and N.D. Daw, A simple model for learning in volatile environments. PLOS Computational Biology, 2020. 16(7): p. e1007963.

20. Raio, C.M., et al., Stress attenuates the flexible updating of aversive value. Proc Natl Acad Sci U S A, 2017. 114(42): p. 11241–11246.

21. Putica, A., J. Agathos, and K. Felmingham, How insights from posttraumatic stress disorder can inform treatment of functional neurological disorder. Nature Reviews Psychology, 2025. 4(10): p. 654–668.

22. Dias-Ferreira, E., et al., Chronic stress causes frontostriatal reorganization and affects decision-making. Science, 2009. 325(5940): p. 621–5.

23. Soares, J.M., et al., Stress-induced changes in human decision-making are reversible. Translational Psychiatry, 2012. 2(7): p. e131–e131.

24. Guitart-Masip, M., et al., Anxiety associated with perceived uncontrollable stress enhances expectations of environmental volatility and impairs reward learning. Sci Rep, 2023. 13(1): p. 18451.

25. Voulgaropoulou, S.D., et al., Perceived chronic stress and impulsivity are associated with reduced learning about the costs and benefits of actions. Learning and Motivation, 2023. 83: p. 101896.

26. Petrowski, K., et al., Norm values and psychometric properties of the short version of the Trier Inventory for Chronic Stress (TICS) in a representative German sample. PLoS One, 2019. 14(11): p. e0222277.

27. Beck, A.T., et al., Comparison of Beck Depression Inventories -IA and -II in psychiatric outpatients. J Pers Assess, 1996. 67(3): p. 588–97.

28. Beck, A.T., et al., An inventory for measuring clinical anxiety: psychometric properties. J Consult Clin Psychol, 1988. 56(6): p. 893–7.

29. Buhr, K. and M.J. Dugas, The intolerance of uncertainty scale: psychometric properties of the English version. Behaviour Research and Therapy, 2002. 40(8): p. 931–945.

30. Rajput, D., et al., Predictability and controllability shape aversive learning and stress responses through independent computational mechanisms. bioRxiv, 2026: p. 2026.08.26.745064.

31. Gracely, R.H., et al., A multiple random staircase method of psychophysical pain assessment. Pain, 1988. 32(1): p. 55–63.

32. Grupe, D.W. and J.B. Nitschke, Uncertainty and anticipation in anxiety: an integrated neurobiological and psychological perspective. Nat Rev Neurosci, 2013. 14(7): p. 488–501.

33. Peters, A., B.S. McEwen, and K. Friston, Uncertainty and stress: Why it causes diseases and how it is mastered by the brain. Progress in Neurobiology, 2017. 156: p. 164–188.

34. Liu, Q., et al., Impact of Chronic Stress on Attention Control: Evidence from Behavioral and Event-Related Potential Analyses. Neuroscience Bulletin, 2020. 36(11): p. 1395–1410.

35. Portes, B., et al., The effects of perceived chronic stress on the fMRI correlates of attentional control in women managers. Arch Womens Ment Health, 2019. 22(3): p. 375–381.

36. Ligneul, R., et al., Stress-sensitive inference of task controllability. Nat Hum Behav, 2022. 6(6): p. 812–822.

37. Wolff, M., et al., Chronic stress, executive functioning, and real-life self-control: An experience sampling study. J Pers, 2021. 89(3): p. 402–421.

38. Havranek, M.M., et al., Uncontrollable and unpredictable stress interacts with subclinical depression and anxiety scores in determining anxiety response. Stress, 2016. 19(1): p. 53–62.

39. Tafet, G.E. and T. Ortiz Alonso, Learned helplessness and learned controllability: from neurobiology to cognitive, emotional and behavioral neurosciences. Frontiers in Psychiatry, 2025. **Volume** 16 **-** 2025.

40. Hoehn-Saric, R. and D.R. McLeod, Anxiety and arousal: physiological changes and their perception. J Affect Disord, 2000. 61(3): p. 217–24.

41. Hu, N., et al., Neural and behavioral dynamics of error processing under chronic stress in healthy young adults. International Journal of Clinical and Health Psychology, 2025. 25(1): p. 100561.

42. Wu, J., et al., The relationship between chronic perceived stress and error processing: evidence from event-related potentials. Sci Rep, 2019. 9(1): p. 11605.

43. S., Albani, C., & Brähler, E. (2012). Factor structure and psychometric properties of the trier inventory for chronic stress (TICS) in a representative German sample. BMC Med Res Methodol, 12, 42.

