## Supplementary File for "Chronic stress reduces behavioural flexibility and autonomic adaptation when aversive outcomes are controllable"

### Supplemental Methods

#### Participants and questionnaires

The TICS-9 used for group classification, described in the main manuscript, contains nine items rated from 1 (never) to 5 (very often), yielding total scores from 9 to 45 and assessing chronic stress across domains including occupational demands, social stress, and persistent worry over the preceding 3 months [1]. Before testing, participants completed the Beck Depression Inventory (BDI [2]), Beck Anxiety Inventory (BAI [3]), and 27-item Intolerance of Uncertainty Scale (IUS). The BDI comprises 21 items scored from 0 to 3 and summed to give a total score of 0–63, with higher scores indicating greater depressive symptom severity; scores of 0–13, 14–19, 20–28, and 29–63 indicate minimal, mild, moderate, and severe symptoms, respectively. The BAI comprises 21 items assessing anxiety symptoms experienced during the preceding week, rated from 0 (“not at all”) to 3 (“a lot”) and summed to give a total score of 0–63, with higher scores indicating greater anxiety symptom severity. For the Intolerance of Uncertainty Scale, prospective intolerance of uncertainty (Factor 1) reflected desire for predictability and active certainty seeking, whereas inhibitory intolerance of uncertainty (Factor 2) reflected difficulty thinking or acting under uncertainty [4]. The IUS comprises 27 items rated from 1 (“not at all characteristic of me”) to 5 (“entirely characteristic of me”), with higher scores indicating greater intolerance of uncertainty. Factor 1 comprises 15 whereas Factor 2 comprises 12 items; factor scores were calculated by summing the items loading on each factor, giving possible ranges of 15–75 and 12–60, respectively. The two factor scores were analysed separately in the secondary questionnaire analyses.

#### Task procedure and design

The task was programmed and presented using SuperLab 6 (Cedrus Corporation, San Pedro, California). Written instructions and a brief practice session preceded each experimental session. On each trial, one of two rock-image cues was presented for 300 ms (±50 ms). Participants predicted whether a snake would appear by pressing the left or right arrow key within 1200 ms of cue onset. The selected prediction was displayed for 1000 ms (±200 ms), followed by the outcome for 1500 ms (±200 ms). Each trial ended with a 2000-ms intertrial interval.

Each session comprised 320 trials. Predictability changed across 10 unsignalled blocks of approximately 25-35 trials, with block order counterbalanced across participants. Participants were informed that cue-outcome probabilities could change during the task and might at times appear random. The controllability manipulation and cue-outcome probabilities for the three predictability levels are described in the main manuscript.

#### Shock calibration and stimulation

Electrical stimulation was controlled using customised LabChart code through a PowerLab interface (ADInstruments, Dunedin, New Zealand) and delivered through an electrode positioned adjacent to the first dorsal interosseous muscle of the left hand. Before each session, shock intensity was individually calibrated using a 4-point scale from 1 (low) to 4 (painful) and an established staircase procedure (Gracely et al., 1988). The stimulation level corresponding to the boundary between uncomfortable (3) and painful (4) was used as the high-intensity shock. The relation between high- and low-intensity stimulation levels is reported in the main manuscript.

#### General data analysis

All preprocessing, computational modelling, and statistical analyses were conducted in MATLAB R2021b (MathWorks, Natick, Massachusetts) using built-in functions, custom scripts, and specialised toolboxes. The VKF was implemented using the publicly available MATLAB code accompanying Piray and Daw [5], and random-effects Bayesian model comparison was performed using the VBA toolbox. Across behavioural, skin conductance, and model-based analyses, outliers were identified separately within each experimental condition as values exceeding ±3 standard deviations from the condition mean and were excluded from the corresponding analysis.

#### Behavioural analysis

Trials without a response within 1200 ms were treated as missed and excluded. Prediction accuracy and reaction time were calculated separately for each predictability level within each controllability condition. Participants with missing data for any predictability block type in either session were excluded from the corresponding behavioural analysis. Condition-specific outlier handling followed the procedure described above.

#### Skin conductance preprocessing

Skin conductance was recorded from the nondominant hand using two Ag/AgCl electrodes attached to the intermediate phalanges of the index and 3rd fingers. Data were exported to MATLAB and preprocessed using PsPM version 5.0 together with custom scripts. Potential movement-related artefacts were identified from the second derivative of the signal; samples with absolute second-derivative values exceeding 3 standard deviations from the participant-specific mean were flagged. Flagged segments and complete recordings were visually inspected to confirm artefacts and identify signal dropout, excessive noise, or unstable baselines. Confirmed artefactual samples were replaced by linear interpolation between adjacent valid samples. Trials were excluded when artefact or signal loss affected more than 30% of samples in the analysis window. Participants were excluded if more than 30% of trials were invalid, if signal amplitude remained below 0.05 µS for more than 20% of the recording, or if baseline drift exceeded 5 µS/min. Eight participants were excluded, leaving 52 for skin conductance analyses. Retained signals were filtered using a first-order Butterworth bandpass filter (0.05-5 Hz), square-root transformed to reduce positive skew, and range-corrected separately for each participant within each condition. Anticipatory skin conductance response was defined as the peak phasic amplitude during the 2 seconds preceding outcome presentation.

#### Volatile Kalman Filter

Trial-wise learning was modelled using the binary Volatile Kalman Filter (VKF; Piray & Daw, 2020), a state-space model designed for environments in which outcome contingencies can change over time. The VKF assumes that observed binary outcomes are generated from an underlying latent state representing the current probability of an outcome. In the binary model, this latent state is expressed in log-odds space and mapped to outcome probability using a logistic sigmoid function. The latent state is allowed to change across trials, while environmental volatility represents the inferred rate at which these underlying outcome contingencies are changing. Beliefs are updated from trial to trial according to prediction error and uncertainty. When inferred volatility is higher, the model increases the learning rate, giving greater weight to recent outcomes and allowing beliefs to adapt more rapidly to changing contingencies. Thus, the model represents learning about both the current outcome probability and how rapidly that probability is changing, rather than treating cue–outcome contingencies as fixed.

The same VKF specification was applied to healthy controls and participants with chronic stress. The model was fitted separately for each participant and session using the full trial sequence to preserve sequential belief updating. Outcomes were coded as $o_{t}$=1 for high-intensity shock and $o_{t}$=0 for low-intensity shock. The fitted model generated trial-wise estimates of surprise, belief uncertainty, environmental volatility, and learning rate. Surprise reflected the absolute discrepancy between the expected and observed outcome, belief uncertainty reflected uncertainty about the current outcome probability, volatility reflected the inferred rate of change in outcome contingencies, and learning rate indexed the extent to which each new outcome updated the current belief. These trial-wise quantities were then used in the subsequent response-model analyses.

Formally, the trial-wise belief about the probability of the observed outcome was represented by a latent mean $m_{t}$ in log-odds space and updated according to

$$m_{t}=m_{t-1}+k_{t}(o_{t}-s(m_{t-1}))$$

where s(.) is the logistic sigmoid and $k_{t}$is the trial-wise learning rate. Model parameters were estimated by maximising the log-likelihood of trial-wise responses using gradient-based optimisation with multiple random initialisations. Posterior estimates were transformed into the VKF parameters lambda, $v_{0}$, corresponding to volatility updating. The fitted model generated trial-wise surprise, calculated as |$o_{t}-s\left( m_{t-1} \right)$|, together with belief uncertainty, volatility, and learning rate. The volatility-update parameter lambda and mean trial-wise learning rate were retained for secondary correlation analyses [5].

#### Response models and Bayesian model comparison

Participant-level general linear models were fitted separately for subjective stress ratings, reaction time, and skin conductance within each controllability condition using ordinary least squares (MATLAB fitlm). For each response measure, three models were estimated, each containing surprise, belief uncertainty, or volatility as the predictor of interest. Predictors were standardised within participant and session. Subjective ratings were modelled as

$$\mathrm{Stress}_{t}=\beta_{0}+\beta_{1}x_{t}+\beta_{2}\mathrm{Stress}_{t-1}+\beta_{3}\mathrm{Accuracy}_{t}+\varepsilon_{t}$$

Reaction times were modelled as

$$\mathrm{RT}_{t}=\beta_{0}+\beta_{1}x_{t}+\beta_{2}\mathrm{Accuracy}_{t}+\varepsilon_{t}$$

Skin conductance responses were modelled as

$\mathrm{SCR}_{t}=\beta_{0}+\beta_{1}x_{t}+\beta_{2}\mathrm{Accuracy}_{t}+\varepsilon_{t}$.

Here, $x_{t}$ denotes surprise, belief uncertainty, or volatility in separate models. This procedure yielded nine response models per participant per session. Models were compared separately by response measure, group, and controllability condition using random-effects Bayesian model comparison. Bayesian Information Criterion values were converted to approximate log model evidence, and exceedance probability was used to quantify the relative evidence for each competing model.

#### Coefficient and correlation analyses

Participant-specific regression coefficients were retained for the complete coefficient displays (Supplementary Fig. S1). For the cross-response analyses shown in Supplementary Fig. S2, coefficients indexing sensitivity to the same computational quantity were correlated across response domains separately by group and controllability condition. Secondary analyses correlated the volatility-update rate (lambda) and mean trial-wise learning rate with prediction accuracy, reaction time, and mean skin conductance response. Questionnaire analyses correlated BDI-II, BAI, and Intolerance of Uncertainty factor scores with task-derived behavioural and model-based measures. Pearson correlations were calculated separately by group, predictability level, and controllability condition.

### Supplementary Results

#### Complete coefficient and cross-response analyses

Complete regression-coefficient distributions from the belief-uncertainty, volatility, and surprise models are shown in Supplementary Fig. S1. Cross-response associations between subjective-rating coefficients and corresponding reaction-time or skin-conductance coefficients are shown in Supplementary Fig. S2. Primary inferential effects from these coefficient analyses are reported in the main manuscript and are not repeated here.

#### Questionnaire and model-derived associations

Beyond the associations reported in the main manuscript, healthy controls showed a positive association between BAI score and mean learning rate under controllable moderate-predictability conditions, r(25) = .47, p = .016. BDI-II score was positively associated with mean learning rate under controllable high-predictability conditions, r(25) = .51, p = .009, and negatively associated with prediction accuracy under controllable moderate-predictability conditions, r(25) = -.41, p = .038. Prospective intolerance of uncertainty (Factor 1) was negatively associated with volatility-update rate under controllable conditions, r(25) = -.41, p = .043, whereas inhibitory intolerance of uncertainty (Factor 2) was positively associated with mean learning rate under controllable high-predictability conditions, r(25) = .44, p = .018. No further significant associations were identified in healthy controls. No additional significant associations beyond those reported in the main manuscript were identified in the chronic stress group.

.

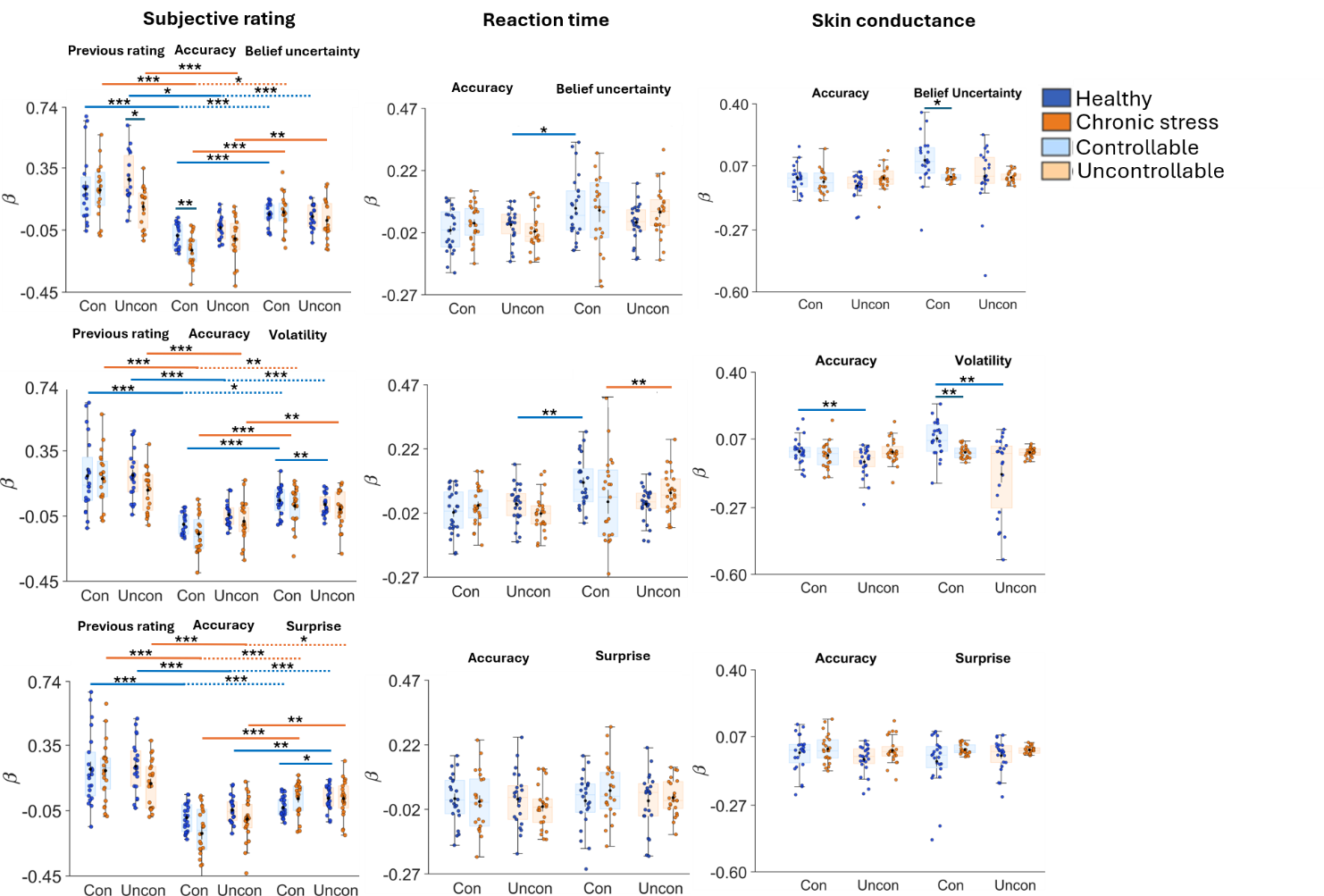

### Supplementary Figures

**Fig. S1. Fig. S1.** Complete regression coefficients for subjective stress ratings, reaction time, and skin conductance across computational models. Regression coefficients (β) are shown for subjective stress ratings (left column), reaction time (middle column), and skin conductance (right column), with rows corresponding to models including belief uncertainty (top), volatility (middle), and surprise (bottom) as the computational predictor. Subjective-rating models include coefficients for the preceding rating, prediction accuracy, and the computational predictor, whereas reaction-time and skin-conductance models include coefficients for prediction accuracy and the computational predictor. Points represent participant-level coefficients for healthy controls and individuals with chronic stress under controllable (Con) and uncontrollable (Uncon) conditions. Blue significance lines indicate within-healthy comparisons, orange lines indicate within-chronic-stress comparisons, and black lines indicate between-group comparisons. For subjective ratings, dotted line segments denote continuations of the corresponding solid lines and are used to distinguish overlapping comparisons. Asterisks indicate statistical significance (p < .05, p < .01, p < .001; Bonferroni-corrected). Selected effects are reported in the main manuscript. Selected effects are reported in the main
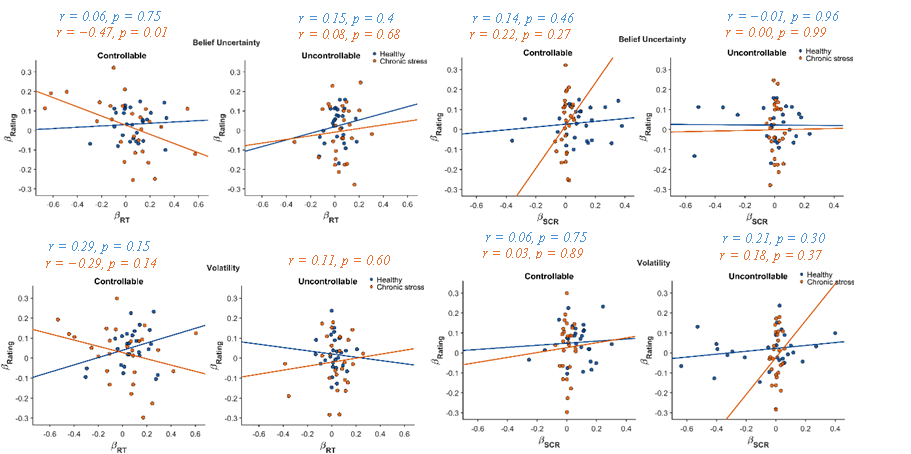
manuscript.

**Fig. S2. Cross-response associations between computational regression coefficients.** Participant-level regression coefficients (β) for subjective stress ratings are plotted against the corresponding reaction-time (RT) and skin-conductance response (SCR) coefficients for belief uncertainty and volatility, separately for controllable and uncontrollable conditions in healthy controls and individuals with chronic stress. Points represent individual participants, and lines indicate least-squares fits. In the chronic stress group, belief-uncertainty coefficients for ratings were negatively associated with those for RT under the controllable condition (r = −0.47, p = 0.01), although this association did not remain significant after Bonferroni correction (Bonf. p = 0.35). No other rating–RT or rating–SCR associations were significant (all p > 0.05).
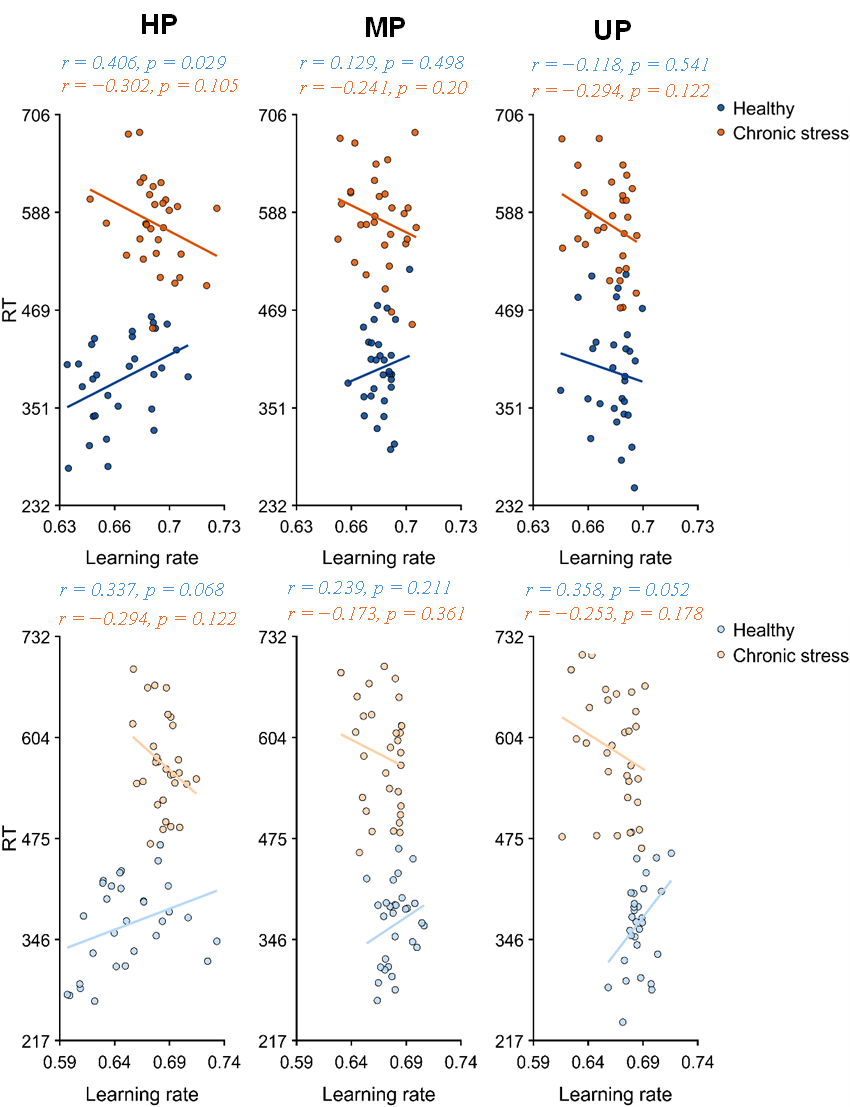

**Fig. S3.** Associations between mean learning rate and reaction time across predictability blocks and controllability conditions. Associations are shown for healthy controls and individuals with chronic stress across highly predictable (HP), moderately predictable (MP), and unpredictable (UP) blocks under controllable and uncontrollable conditions. The upper row shows the controllable condition, and the lower row shows the uncontrollable condition. Healthy controls are shown in blue and individuals with chronic stress in orange. Darker shades indicate the controllable condition, and lighter shades indicate the uncontrollable condition. Points represent individual participants and lines indicate least-squares fits. In healthy controls, higher learning rates were associated with longer reaction times in highly predictable blocks under the controllable condition, r(27) = 0.406, p = 0.029. No other learning-rate–reaction-time associations were significant, all p ≥ 0.052.

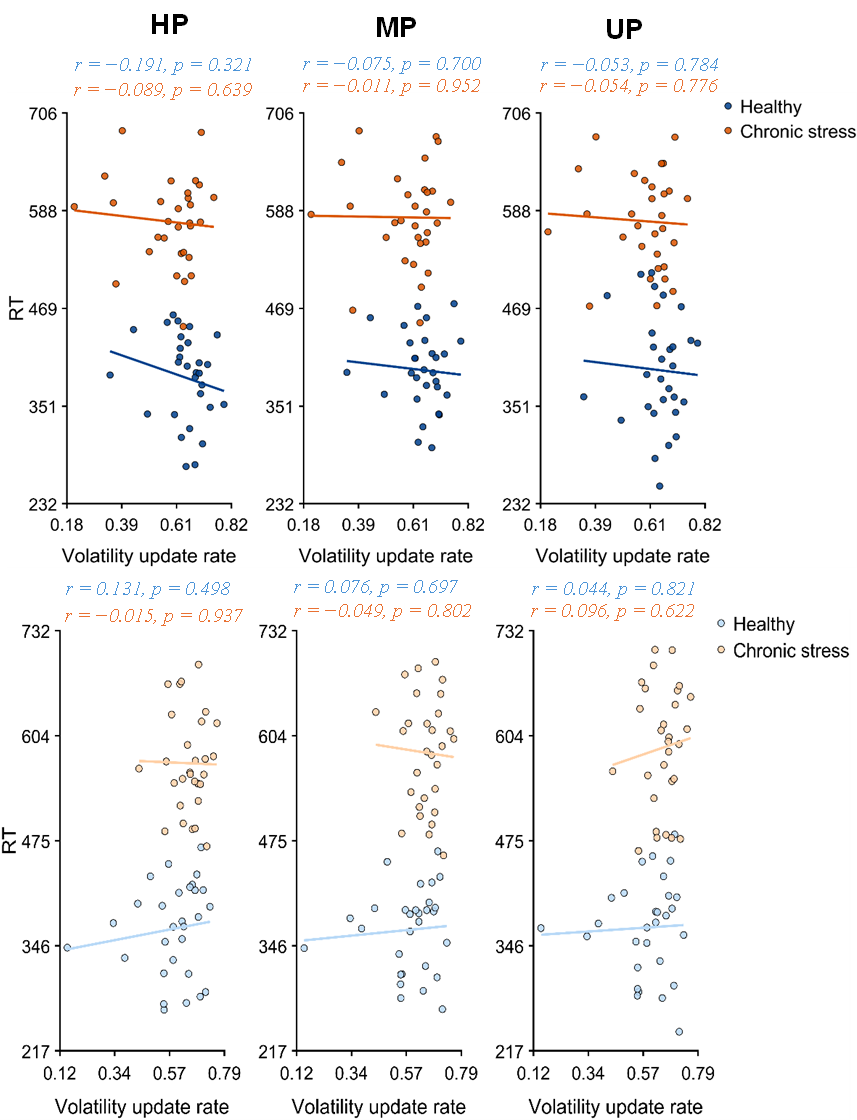

**Fig. S4**. Associations between volatility update rate and reaction time across predictability blocks under controllable and uncontrollable conditions. Associations between volatility update rate and reaction time (RT) are shown for healthy controls and individuals with chronic stress across highly predictable (HP; left), moderately predictable (MP; middle), and unpredictable (UP; right) blocks. The top row shows the controllable condition, and the bottom row shows the uncontrollable condition. Healthy controls are shown in dark blue under the controllable condition and light blue under the uncontrollable condition; individuals with chronic stress are shown in dark orange under the controllable condition and light orange under the uncontrollable condition. Points represent individual participants, and lines indicate least-squares fits. Pearson’s r and corresponding p values are reported above each panel in the colour corresponding to each group and condition. No significant associations between volatility update rate and RT were observed in either group across predictability blocks under either controllability condition (all p ≥ .05).

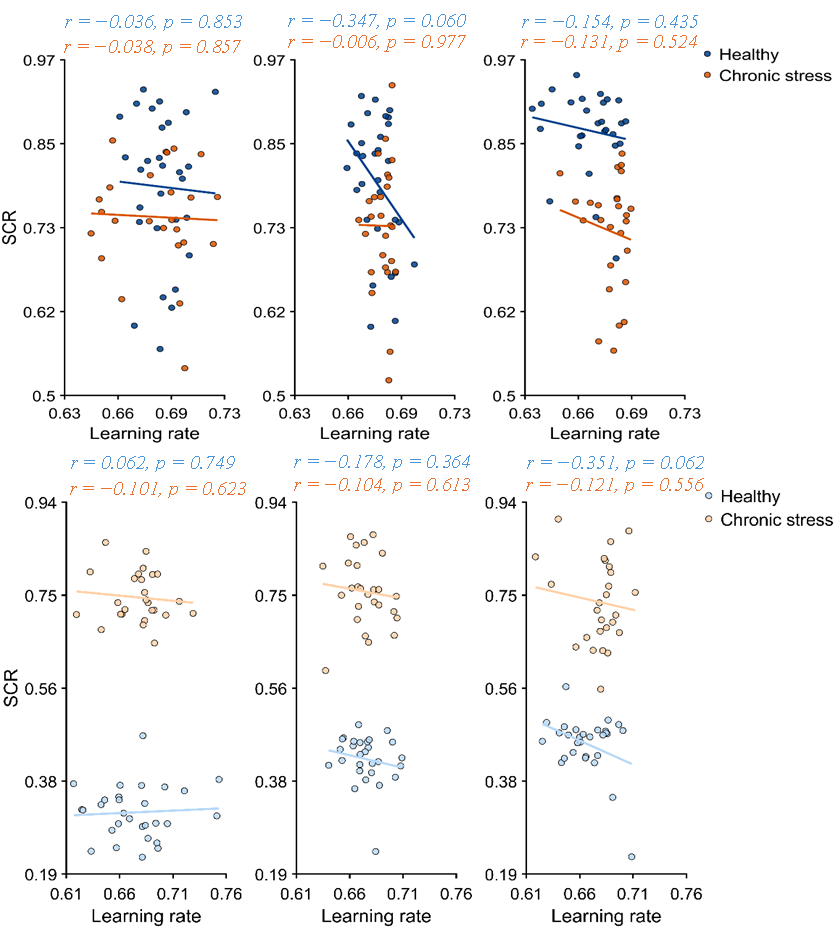

**Fig. S5.** Associations between mean learning rate and skin conductance response across predictability blocks under controllable and uncontrollable conditions. Associations between mean learning rate and skin conductance response (SCR) are shown for healthy controls and individuals with chronic stress across highly predictable (HP; left), moderately predictable (MP; middle), and unpredictable (UP; right) blocks. The top row shows the controllable condition, and the bottom row shows the uncontrollable condition. Healthy controls are shown in dark blue under the controllable condition and light blue under the uncontrollable condition; individuals with chronic stress are shown in dark orange under the controllable condition and light orange under the uncontrollable condition. Points represent individual participants, and lines indicate least-squares fits. Pearson’s r and corresponding p values are reported above each panel in the colour corresponding to each group and condition. No significant associations between learning rate and SCR were observed in either group across predictability blocks under either
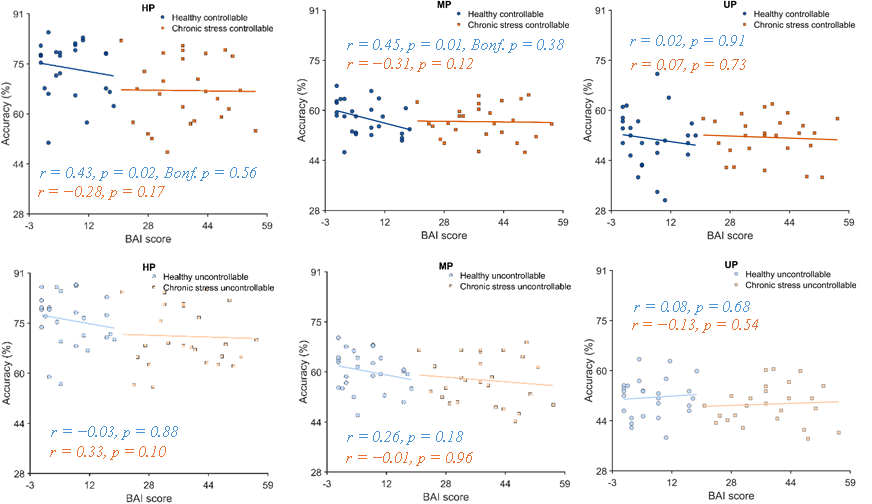
controllability condition (all p ≥ .05).

**Fig. S6.** Associations between Beck Anxiety Inventory (BAI) scores and prediction accuracy across predictability blocks under controllable and uncontrollable conditions. Associations between BAI scores and prediction accuracy are shown for healthy controls and individuals with chronic stress across highly predictable (HP; left), moderately predictable (MP; middle), and unpredictable (UP; right) blocks. The top row shows the controllable condition, and the bottom row shows the uncontrollable condition. Healthy controls are shown in dark blue under the controllable condition and light blue under the uncontrollable condition; individuals with chronic stress are shown in dark orange under the controllable condition and light orange under the uncontrollable condition. Points represent individual participants, and lines indicate least-squares fits. Pearson’s r and corresponding p values are reported for each association, with Bonferroni-corrected p values additionally shown where applicable.

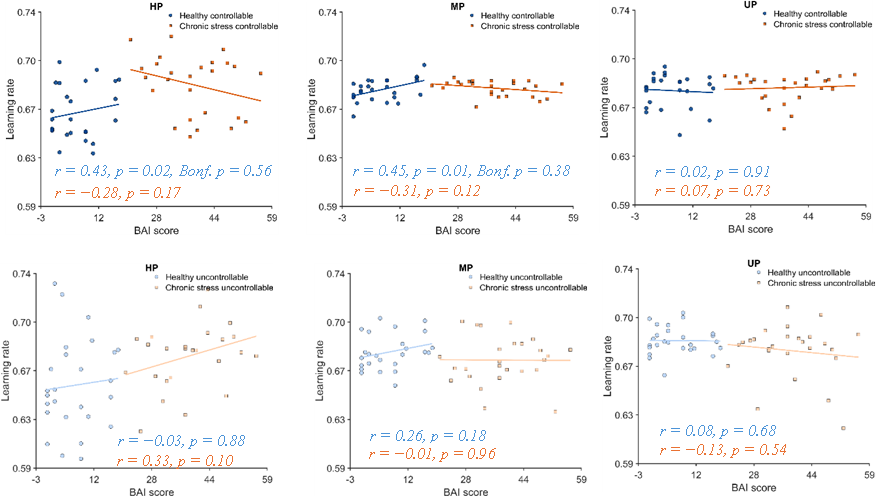

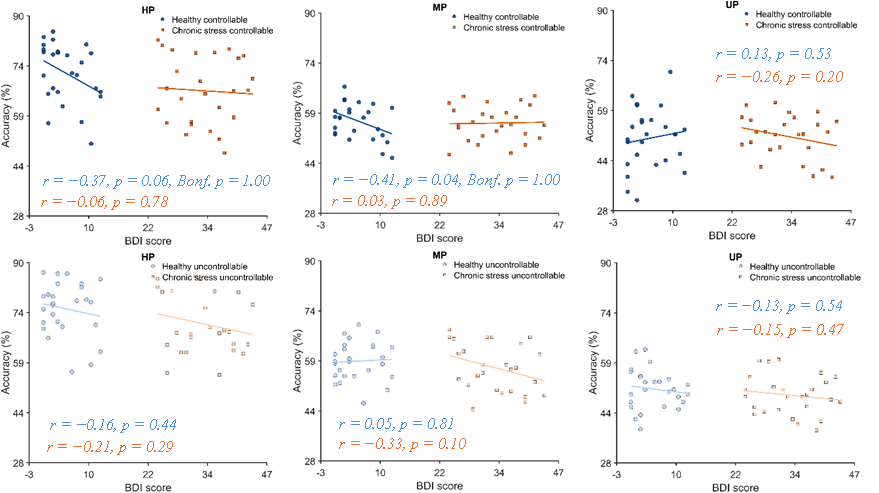
**Fig. S7.** Associations between Beck Anxiety Inventory (BAI) scores and mean learning rate across predictability blocks under controllable and uncontrollable conditions. Associations between BAI scores and mean learning rate are shown for healthy controls and individuals with chronic stress across highly predictable (HP; left), moderately predictable (MP; middle), and unpredictable (UP; right) blocks. The top row shows the controllable condition, and the bottom row shows the uncontrollable condition. Healthy controls are shown in dark blue under the controllable condition and light blue under the uncontrollable condition; individuals with chronic stress are shown in dark orange under the controllable condition and light orange under the uncontrollable condition. Points represent individual participants, and lines indicate least-squares fits. Pearson’s r and corresponding p values are reported for each association, with Bonferroni-corrected p values shown for associations with uncorrected p < .05. No associations remained significant after Bonferroni correction.

**Fig. S8.** Associations between Beck Depression Inventory (BDI) scores and prediction accuracy across predictability blocks under controllable and uncontrollable conditions. Associations between BDI scores and prediction accuracy are shown for healthy controls and individuals with chronic stress across highly predictable (HP; left), moderately predictable (MP; middle), and unpredictable (UP; right) blocks. The top row shows the controllable condition, and the bottom row shows the uncontrollable condition. Healthy controls are shown in dark blue under the controllable condition and light blue under the uncontrollable condition; individuals with chronic stress are shown in dark orange under the controllable condition and light orange under the uncontrollable condition. Points represent individual participants, and lines indicate least-squares fits. Pearson’s r and corresponding p values are reported for each association, with Bonferroni-corrected p values additionally shown where applicable. In healthy controls, higher BDI scores were associated with lower prediction accuracy in the MP block under the controllable condition (r = −0.41, p = .04), but this association did not remain significant after
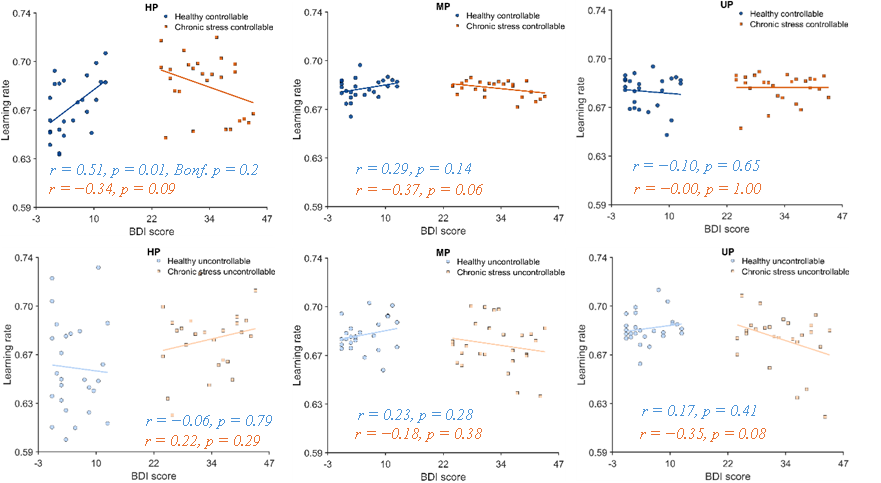
Bonferroni correction (Bonf. p = 1.00). No other associations were significant (all p > .05).

**Fig. S9.** Associations between Beck Depression Inventory (BDI) scores and mean learning rate across predictability blocks under controllable and uncontrollable conditions. Associations between BDI scores and mean learning rate are shown for healthy controls and individuals with chronic stress across highly predictable (HP; left), moderately predictable (MP; middle), and unpredictable (UP; right) blocks. The top row shows the controllable condition, and the bottom row shows the uncontrollable condition. Healthy controls are shown in dark blue under the controllable condition and light blue under the uncontrollable condition; individuals with chronic stress are shown in dark orange under the controllable condition and light orange under the uncontrollable condition. Points represent individual participants, and lines indicate least-squares fits. Pearson’s r and corresponding p values are reported for each association, with Bonferroni-corrected p values additionally shown where applicable. In healthy controls, higher BDI scores were associated with higher mean learning rate in the HP block under the controllable condition (r = 0.51, p = .01), but this association did not remain significant after
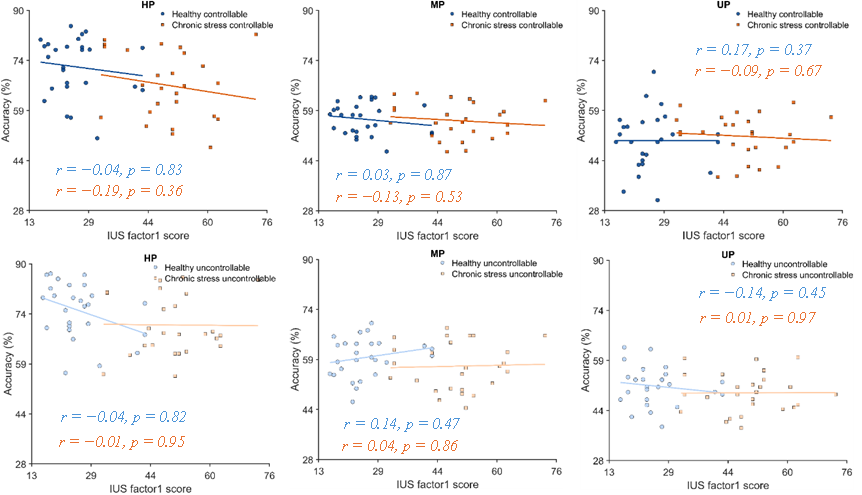
Bonferroni correction. No other associations were significant (all p > .05).

**Fig. S10.** Associations between prospective intolerance of uncertainty (IUS Factor 1) scores and prediction accuracy across predictability blocks under controllable and uncontrollable conditions. Associations between IUS Factor 1 scores and prediction accuracy are shown for healthy controls and individuals with chronic stress across highly predictable (HP; left), moderately predictable (MP; middle), and unpredictable (UP; right) blocks. The top row shows the controllable condition, and the bottom row shows the uncontrollable condition. Healthy controls are shown in dark blue under the controllable condition and light blue under the uncontrollable condition; individuals with chronic stress are shown in dark orange under the controllable condition and light orange under the uncontrollable condition. Points represent individual participants, and lines indicate least-squares fits. Pearson’s r and corresponding p values are reported for each association. No significant associations between IUS Factor 1 scores and prediction accuracy were observed in either group across predictability blocks under
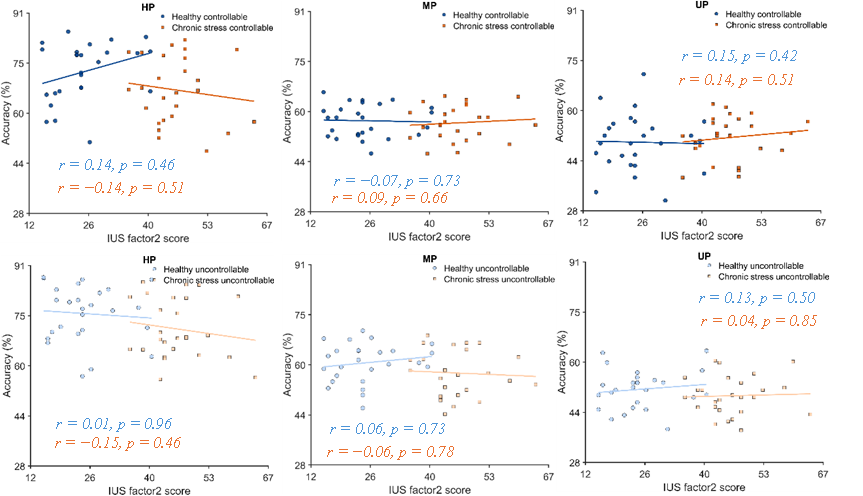
either controllability condition (all p > .05).

**Fig. S11.** Associations between inhibitory intolerance of uncertainty (IUS Factor 2) scores and prediction accuracy across predictability blocks under controllable and uncontrollable conditions. Associations between IUS Factor 2 scores and prediction accuracy are shown for healthy controls and individuals with chronic stress across highly predictable (HP; left), moderately predictable (MP; middle), and unpredictable (UP; right) blocks. The top row shows the controllable condition, and the bottom row shows the uncontrollable condition. Healthy controls are shown in dark blue under the controllable condition and light blue under the uncontrollable condition; individuals with chronic stress are shown in dark orange under the controllable condition and light orange under the uncontrollable condition. Points represent individual participants, and lines indicate least-squares fits. Pearson’s r and corresponding p values are reported for each association. No significant associations between IUS Factor 2 scores and prediction accuracy were observed in either group across predictability blocks under
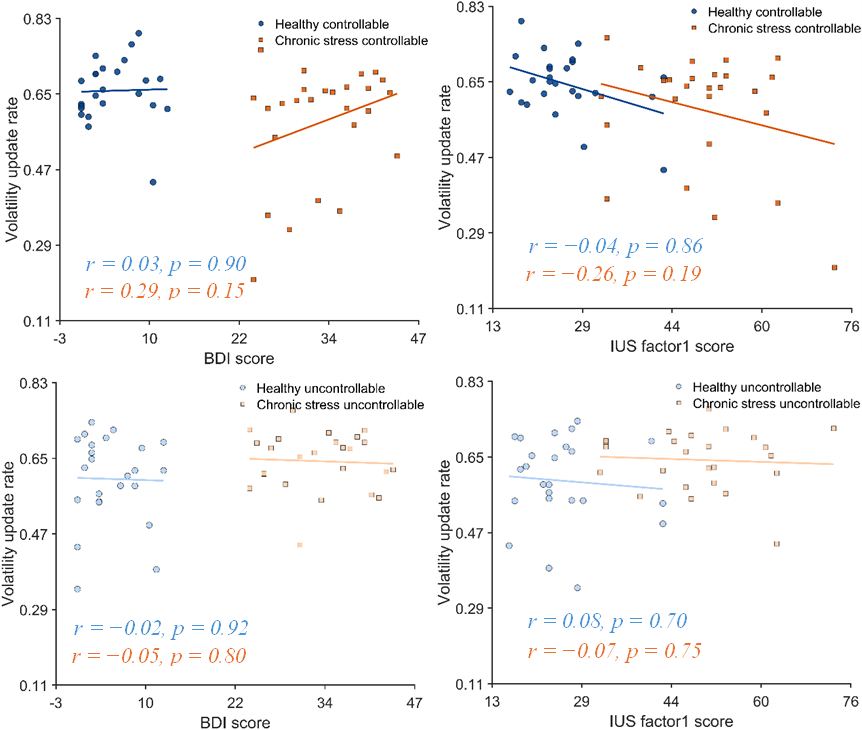
either controllability condition (all p > .05).

**Fig. S12.** Associations of Beck Depression Inventory-II (BDI-II) and prospective intolerance of uncertainty (IUS Factor 1) scores with volatility update rate under controllable and uncontrollable conditions. Associations between BDI-II scores (left) and IUS Factor 1 scores (right) and volatility update rate are shown for healthy controls and individuals with chronic stress. The top row shows the controllable condition, and the bottom row shows the uncontrollable condition. Healthy controls are shown in dark blue under the controllable condition and light blue under the uncontrollable condition; individuals with chronic stress are shown in dark orange under the controllable condition and light orange under the uncontrollable condition. Points represent individual participants, and lines indicate least-squares fits. Pearson’s r and corresponding p values are reported for each association. No significant associations between BDI-II or IUS Factor 1 scores and volatility update rate were observed in either group under either controllability condition (all p > .05).

### Supplemental References

1.	Petrowski, K., et al., *Norm values and psychometric properties of the short version of the Trier Inventory for Chronic Stress (TICS) in a representative German sample.* PLoS One, 2019. **14**(11): p. e0222277.

2.	Beck, A.T., et al., *Comparison of Beck Depression Inventories -IA and -II in psychiatric outpatients.* J Pers Assess, 1996. **67**(3): p. 588-97.

3.	Beck, A.T., et al., *An inventory for measuring clinical anxiety: psychometric properties.* J Consult Clin Psychol, 1988. **56**(6): p. 893-7.

4.	Buhr, K. and M.J. Dugas, *The intolerance of uncertainty scale: psychometric properties of the English version.* Behaviour Research and Therapy, 2002. **40**(8): p. 931-945.

5.	Piray, P. and N.D. Daw, *A simple model for learning in volatile environments.* PLOS Computational Biology, 2020. **16**(7): p. e1007963.
